# Keratin 15 promotes a progenitor cell state in basal keratinocytes of skin epidermis

**DOI:** 10.1101/2025.02.27.640633

**Authors:** Catherine J. Redmond, Sarah N. Steiner, Erez Cohen, Craig N. Johnson, Nurhan Özlü, Pierre A. Coulombe

## Abstract

The type I intermediate filament proteins keratin 14 (K14) and keratin 15 (K15) are common to all complex epithelia. K14 is highly expressed by progenitor keratinocytes, in which it provides essential mechanical integrity and gates keratinocyte entry into differentiation by sequestering YAP1, a transcriptional effector of Hippo signaling, to the cytoplasm. K15 has long been used as a marker of hair bulge stem cells, though its specific role in skin epithelia is unknown. Here we show that the lack of two biochemical determinants, a cysteine residue within the stutter motif of the central rod domain and a 14-3-3 binding site in the N-terminal head domain, renders K15 unable to effectively sequester YAP1 in the cytoplasm like K14 does. We combine insight obtained from cell culture and transgenic mouse models with computational analyses of transcriptomics data and propose a model in which a higher K15:K14 ratio promotes a progenitor state and antagonizes differentiation in keratinocytes of the epidermis.

**SUMMARY:** The type I intermediate filament keratin 15 (K15) exhibits preferential expression in a subset of skin keratinocytes with stem-like properties. We report that K15 promotes the progenitor state by counteracting the ability of the related K14 to negatively regulate YAP1, a terminal effector of Hippo signaling.

## INTRODUCTION

Complex epithelia function as physical, chemical and immunological barriers in many organs and tissues across the body, including the skin, eye, oral mucosa, and reproductive tract. The barrier function of these epithelia is maintained throughout life by carefully balancing the growth, via proliferation, and loss, via differentiation, of keratinocytes, the predominant cell type within complex epithelia, through the process of epithelial homeostasis (Blanpain and Fuchs, 2009; Wells and Watt, 2018).

Keratinocytes can be characterized by their denominal keratin intermediate filament (IF) proteins. Owing to their diversity (N=54 in humans), differentiation-dependent regulation, unique mechanical properties, and post-translational modifications, keratin IF proteins play important roles in specific aspects of keratinocyte biology in addition to maintenance of epithelial integrity (Cohen et al., 2022; Fuchs, 1995; Schweizer et al., 2006). The transcriptional and post-translational regulation of keratin genes and proteins, and their impact on the complex process of epithelial homeostasis, are under active investigation (Li et al., 2023). In order to sustain the long-term regeneration of complex epithelia, epithelial homeostasis requires stem-like progenitor keratinocytes that maintain mitotic competency over decades of life (Blanpain and Fuchs, 2009; Wells and Watt, 2018). Here, we provide evidence for a mechanism that prevents the differentiation-related premature loss of progenitor keratinocytes through the regulation of the mechanosensitive transcriptional co-activator YAP1 by keratin IF proteins.

Common to all complex, stratified (e.g., epidermis) and pseudostratified epithelia (e.g., trachea) is the expression of the type I keratins 14 and 15 (K14, K15) and their type II co-polymerization partner K5 (Fuchs, 1995; Schweizer et al, 2006). K5 and K14 mRNA and proteins are prominently co-expressed in all keratinocytes residing in the basal layer of stratified epithelia (Cohen et al., 2022). K5/K14 IFs comprise the majority of the mechanical resilience of these cells, protecting them against shear- and trauma-induced lysis (Coulombe et al., 2009; Ramms et al., 2013). In support of the latter, small, dominantly-acting missense variants affecting the polymerization and mechanical properties of either K5 or K14 are causative for epidermolysis bullosa simplex (EBS), a rare genetic skin disorder in which basal layer keratinocytes are fragile and shear or lyse in response to trauma (Bonifas et al., 1991; Coulombe et al., 1991; Coulombe et al., 2009). In striking contrast, very little is known about K15, other that the intriguing attribute that it is correlated, cell-autonomously, with stemness in the skin (Bose et al., 2013; Lyle et al., 1998) and other stratified epithelia (Giroux et al., 2017).

Additional to its well-defined structural function, a novel role for K14 recently came to the fore through the characterization of mice carrying a *Krt14* C373A mutant allele (Guo et al., 2020). *Krt14* C373A mutant mice were generated to study the physiological significance of a disulfide bond discovered in a crystal structure of the interacting 2B coiled-coil rod domains of human K5 and K14 (Lee et al., 2012). This homotypic transdimer disulfide linkage was shown to involve cysteine (C) 367, located in the 2B domain of human K14 (Lee et al., 2012) (note: C367 in human K14 corresponds to C373 in mouse K14). What distinguishes C367 in human and C373 in mouse K14 from other cysteines is its location in the second residue of the so-called stutter motif, a four-residue disruption of the long-range heptad repeats that is nearly perfectly conserved across the entire superfamily of IF proteins (Lee et al., 2012). Intriguingly, cysteine residues occur in the second position of the stutter in several type I keratins expressed in surface epithelia, including the differentiation-specific K9 and K10 (Lee et al., 2012) and the stress-induced K16 and K17 (Guo et al., 2020). A subsequent study reporting on the atomic structure of the interacting 2B domains of K10 and its partner K1 also uncovered a homotypic disulfide linkage mediated by the K10 stutter cysteine (Bunick and Milstone, 2017).

Live imaging studies of skin keratinocytes in primary culture revealed that the K14 stutter cysteine is required for the establishment of a stable perinuclear filament network (Feng and Coulombe, 2015b), a trademark feature in early-stage differentiating keratinocytes of surface epithelia (Coulombe et al., 1989; Lee et al., 2012). A *Krt14* knock-in mutant C373A mouse was generated to test the physiological role of C373-dependent disulfide bonding. Loss of K14 C373 in these mutant mice caused a delay in the initiation of keratinocyte differentiation and aberrant terminal differentiation, ultimately resulting in a barrier defect (Guo et al., 2020). Assessment of K14 cysteine mutant properties in newborn skin keratinocytes in primary culture coupled with a proteomics screen for K14-interacting proteins revealed that, unlike its wildtype counterpart, the K14 C373A protein is unable to promote the cytoplasmic retention of YAP1, a co-transcriptional effector of Hippo signaling (Totaro et al., 2018), as epidermal keratinocytes initiate differentiation, both *in vivo* and in culture (Guo et al., 2020). Alongside YAP1 dysregulation, the scaffolding protein stratifin/14-3-3σ, a known YAP1-interacting protein (Schlegelmilch et al., 2011), occurs in an aggregated rather than the normal diffuse pattern in suprabasal epidermal keratinocytes of *Krt14* C373A mice (Guo et al., 2020). These findings guided the development of a model whereby the stutter cysteines in K14 and K10 are required for the cytoplasmic retention of YAP1, in a 14-3-3-dependent manner, thus mediating the initiation (via K14) and sustainment of differentiation (via K10) in epidermal keratinocytes.

Relative to K14, the K15 mRNA and protein are expressed in a subset of keratinocytes in the basal layer (Jonkman et al., 1996; Waseem et al., 1999). Studies that followed up on the discovery of K15’s enrichment in the hair bulge (Lyle et al., 1998) strengthened the correlation involving K15 and epithelial stem cells of the skin (Bose et al., 2013). K15 has since been described as a marker for progenitor cells in several epithelial-rich organs and its expression has been correlated to the development and severity of cancers in these tissues (Bose et al., 2013). Despite the breadth of work relating K15 to stemness and cancer, a functional role has yet to be assigned to this type I keratin. In human and mice lacking K14 protein, K15 and K5 form fine (“wispy”) filaments in basal keratinocytes that fail to compensate for the essential structural function of K14/K5 filaments (Chan et al., 1994; Jonkman et al., 1996; Lloyd et al., 1995; Rugg et al., 1994). Intriguingly, unlike K14 or K10, K15 does not feature a cysteine at the second position of the stutter motif in its 2B domain.

Here, we combine biochemical, cell biological, histological, computational and physiological evidence to present evidence that, through its spatially unique distribution and expression levels in epidermis, its lack of a stutter cysteine and therefore a lesser ability to promote the cytoplasmic sequestration of YAP1, K15 functions as a negative regulator of keratinocyte differentiation and promotes the progenitor cellular state in skin epithelia.

## RESULTS

The location of two key biochemical determinants in K14 protein, the “SCRAPS” motif in the N-terminal head domain and a cysteine residue located in the stutter motif of coil 2 in the central α-helical rod domain, and the corresponding sequences in K15, are depicted in Fig. 1A.

**Figure 1.**
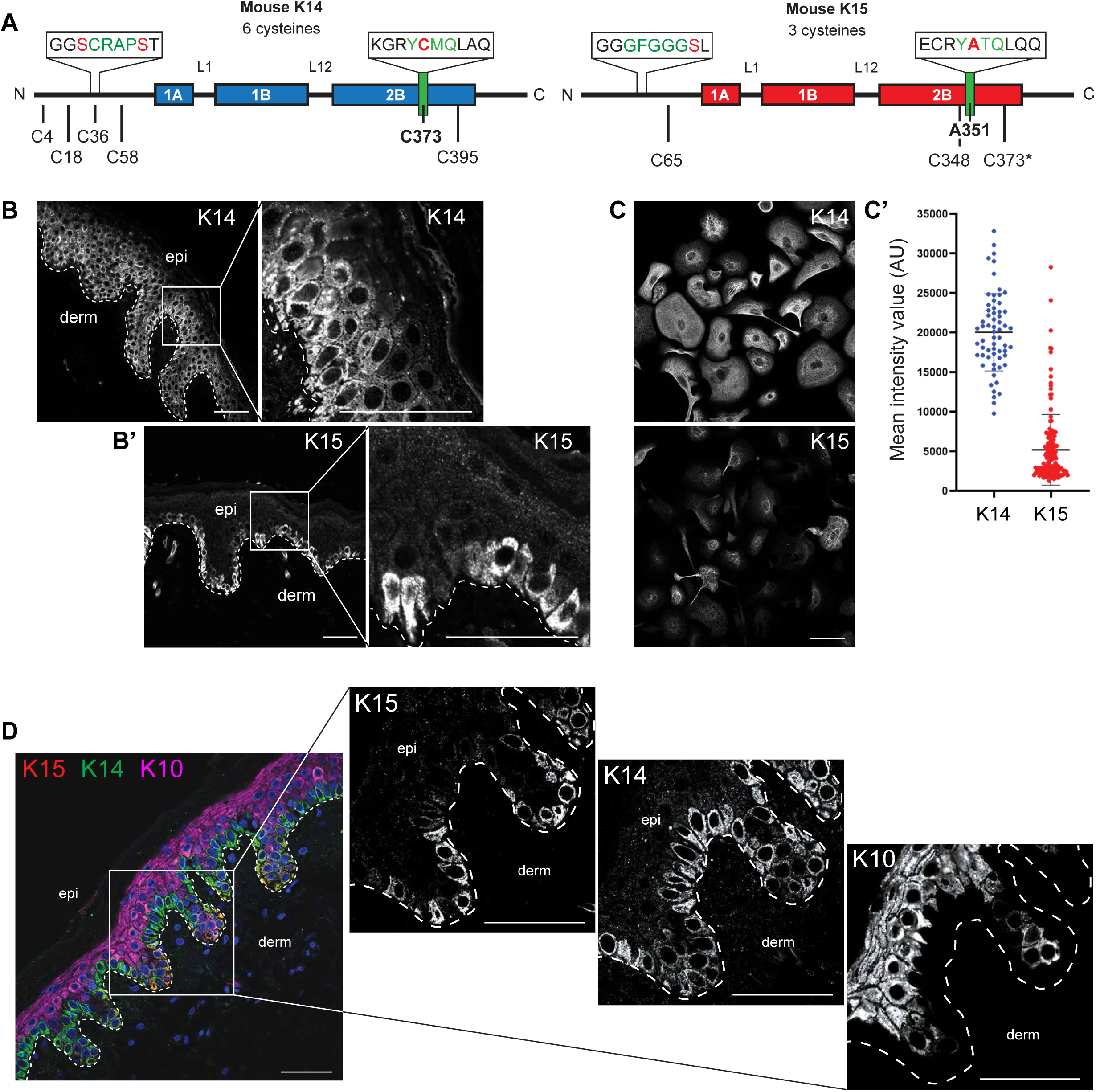
K14 and K15 display distinct staining patterns in vivo and ex vivo. **A.** Diagram of K14 and K15 primary structures. Boxes represent coiled-coil regions. Schematic comparing the orthologous residues between K14 and K15 in the putative regulatory regions in the head and 2B rod domains. Mouse K14 contains 4 cysteines in the head domain and 2 in the 2B rod domain. Mouse K15 contains 1 cysteine residue in the head domain and 2 in the 2B rod domain. K15 does not have a cysteine orthologous to mK14 C373 (mK15 C373 is downstream of the mK15 stutter region). **B.** Individual immunostaining with magnified inset for keratin 14 (K14) in formalin-fixed, paraffin embedded human buttocks skin. K14 individual immunostaining extends into the suprabasal compartment. Dashed line specifies the basal lamina; epi, epidermis; derm, dermis. Scale bar, 50 µm. **B’.** Individual immunostaining with magnified inset for keratin 15 (K15) in formalin-fixed, paraffin embedded human buttocks skin. K15 individual immunostaining is restricted to the basal layer. Dashed line specifies the basal lamina; epi, epidermis; derm, dermis. Scale bar, 50 µm. **C.** Individual immunostaining for keratin 14 (K14) and keratin 15 (K15) in skin keratinocytes harvested from neonatal *Krt14^C373A/WT^*pups and seeded in primary culture. Scale bar, 50 µm. **C’.** Scatter plots representing the K14 and K15 mean intensity values of individually traced cells in Fig. 1C (mean ± SD). **D.** Multiplexed immunostaining for keratin 15 (K15, red), keratin 14 (K14, green), and keratin 10 (K10, magenta). Nuclei are counterstained with DAPI (blue). Dashed line specifies the basal lamina. epi, epidermis; derm, dermis. Scale bar, 50 µm.

### K15 occurs in a spatially defined pool of progenitor keratinocytes in human epidermis

K14 is widely considered, and used, as a marker for “mitotically-competent progenitor keratinocytes” in the basal layer of stratified epithelia, including the epidermis. However, several studies have shown that the expression pattern of K14, at both the protein (Fuchs and Green, 1980) and mRNA levels (Stoler et al., 1988), is broader and includes early-stage differentiating keratinocytes in the lower suprabasal layers of epidermis (Cohen et al., 2022). Immunostaining for K14 in human thin (buttock) skin sections corroborates this (Fig. 1B). K15 displays a distinct immunostaining pattern, in that it is largely excluded from the suprabasal compartment and shows markedly brighter signal in basal keratinocytes located in bottom of rete ridges (Fig. 1B’). Analyses of newborn mouse skin keratinocytes in primary culture under calcium-free, growth-promoting conditions shows that bright K14 immunostaining occurs in all keratinocytes whereas K15 is only present in a subset of the cells (Fig. 1C; quantitation shown in Fig. 1C’), consistent with observations made in human skin tissue sections. Co-staining for K15, K14, and K10 in thin epidermis shows a strict exclusion between the K15^+^ basal and K10^+^ suprabasal compartments and K14^+^ cells showing a gradation between the two (Fig. 1D). These findings extend previous studies by several groups (Jonkman et al., 1996; Porter et al., 2000; Waseem et al., 1999; Webb et al., 2004; Whitbread and Powell, 1998; Zhan et al., 2007) and establish that the striking differences in K14 and K15 expression patterns are largely conserved in human and mouse.

### Single cell transcriptomics reveal attributes that uniquely define KRT15-expressing keratinocytes

The availability of single cell (sc) RNA-seq datasets affords an opportunity to gain insight into the significance of keratin gene expression in human skin (Cohen et al., 2022). In healthy abdominal (thin) skin (Suppl. Fig. 1A), the majority of keratinocytes can be partitioned into two subsets that exhibit either a *KRT14*-high or *KRT10*-high signature, representing progenitor or differentiating status, respectively (Suppl. Fig. 1B,C; see Cohen et al., 2022). A third, sizable group of keratinocytes exhibit a hybrid character with appreciable expression of both *KRT14* and *KRT10* (Suppl. Fig. 1D) (Cohen et al., 2022). By comparison, *KRT15* expression (Suppl. Fig. 1E) occurs only in a subset of *KRT14-*expressing keratinocytes (Fig. 2A). These *KRT15*-expressing cells show minimal overlap with *KRT10*-expressing cells (Fig. 2A). Directly relating *KRT15* to *KRT14* reads across the population of individual keratinocytes in human trunk skin (N=26,063 cells) (Cohen et al., 2022), in the form of a scatter plot, further highlights that *KRT15*-expressing keratinocytes are a subset of *KRT14*-expressing ones (Fig. 2B). Additionally, we find that the number of *KRT15* reads per cell is, overall, significantly lower than that of *KRT14* in trunk skin keratinocytes (Fig. 2C). These findings are consistent with the immunostaining data reported in Fig. 1.

**Figure 2.**
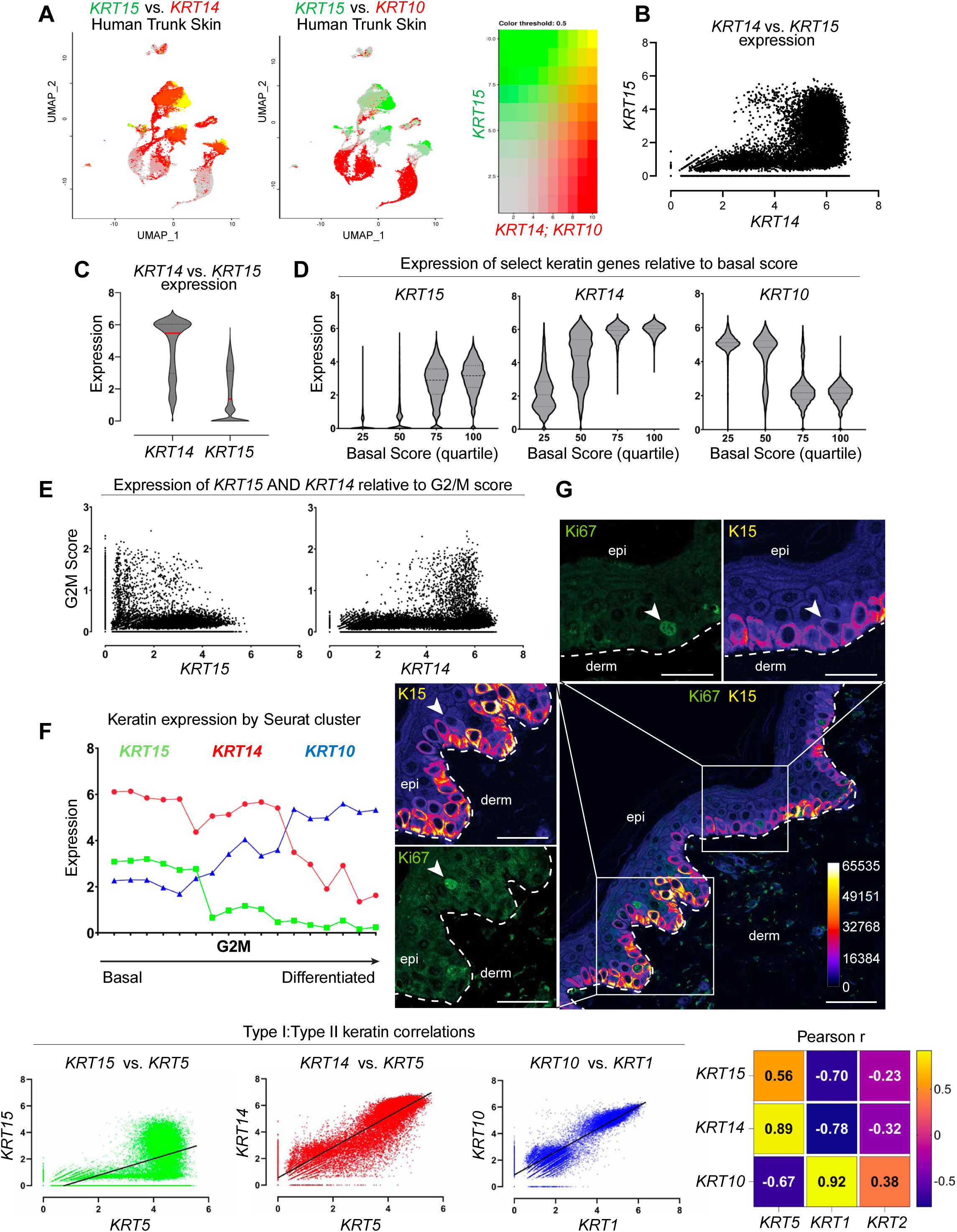
Relationships between *KRT14, KRT15 and KRT10* in human trunk skin. **A.** UMAP of human trunk epidermis highlighting *KRT15* levels (green) and *KRT14* or *KRT10* levels (red) across Seurat clusters. **B.** Single cell expression of *KRT15* mapped against *KRT14* levels in 26,063 keratinocytes in the human trunk epidermis dataset. **C.** Distribution of *KRT14* and *KRT15* expression across all keratinocytes in dataset. **D.** Distribution of *KRT15, KRT14* and *KRT10* expression in cells exhibiting increasing levels of ‘Basal Score’ signature genes (see “Methods” for definition of composite scores). **E.** Expression of G2M composite score relative to *KRT14* and *KRT15* levels across keratinocytes in human trunk epidermis. **F.** Analysis of average keratin expression level across 17 Seurat clusters matching keratinocyte signatures in the human trunk epidermis dataset. *KRT14* (red, circle), *KRT15* (green, square) and *KRT10* (blue, triangle) assessed per cluster. Clusters are sorted on the X axis according to the highest basal score to differentiation score ratio. The Seurat cluster showing the highest G2M composite score, on average, is highlighted. **G.** Indirect immunofluorescence with magnified insets of Ki67 and K15 in formalin-fixed, paraffin embedded human buttocks skin. K15 channel has been false-colored with the “Fire” lookup table to show signal intensity. Strongly Ki67-staining nuclei are frequently K15-low basal keratinocytes. Calibration bar displays the color coding for their corresponding K15 Mean Gray Value intensities. Scale bar, 50 µm. **H.** Comparison of linear pairwise expression of type I and type II keratins *KRT15/KRT5* (green, R^2^=0.31), *KRT14/KRT5* (red, R^2^=0.79), *KRT10/KRT1* (blue, R^2^=0.84) across all keratinocytes in the human trunk epidermis dataset. **H’.** Heatmap summarizing Pearson (r) correlations between type I and type II keratins across the dataset.

To probe deeper into the transcriptional signature that reflects the progenitor cellular state, we next performed a supervised analysis of the entire keratinocyte population of thin epidermis according to composite scores (Cohen et al., 2022; Cohen et al., 2024): a “basal score” to identify cells with high transcriptional activity for genes relating to the extracellular matrix (and thus likely to reside in the basal compartment), a “differentiation score” to identify cells engaged into differentiation (e.g., cell-cell adhesion components), and “G2/M” and “S” scores to identify cells undergoing mitosis. These scores are formulated independently of any keratin gene (see Methods and Cohen et al., 2022).

First, we sorted individual keratinocytes according to their expression of a given keratin gene as a function the basal score, with the latter binned into quartiles (percentiles 0-25, 26-50, 51-75 and 76-100). This analysis revealed that *KRT15* expression shows a clear bias towards keratinocytes with a high basal score, *KRT10* expression shows a clear bias towards keratinocytes with a low basal score, while *KRT14* expression shows a mixed identity (Fig. 2D).

Second, we related keratin expression to the G2/M composite score, which reflects engagement in mitosis, via a scatterplot. This analysis showed that even though they exhibit a high basal score, high *KRT15*-expressing keratinocytes show very low G2M scores, unlike high *KRT14*-expressing cells (Fig. 2E). The same conclusion is attained when relating *KRT15* and *KRT14* expression to a composite score for S Phase (Suppl. Fig. 1F,G). By contrast, the relationship between *KRT10* expression levels and the G2/M score is clearly bimodal (Suppl. Fig. 1H), consistent with the recent demonstration of a transitional population of epidermal keratinocytes that have initiated differentiation and yet are mitotically-active (Cockburn et al., 2022; Cohen et al., 2022; Lin et al., 2020).

Third, we rank-ordered the Seurat clusters (Suppl. Fig. 1A) according to the ratio of their “basal” and “differentiation” scores. This analysis revealed a clear sequence whereby *KRT15* occurs in cells with a clear basal identity, *KRT14* occurs across cells with either a basal or a differentiating identity, and *KRT10* expression is clearly polarized to differentiating cells (Fig. 2F). Interestingly, the Seurat cluster enriched for G2/M keratinocytes maps to cells that feature an intermediate basal-to-differentiation score in the distribution reported in Fig. 2F.

To further substantiate the exclusion of *KRT15*-expressing cells from the subpopulation of actively mitotic G2M cells, we next performed co-immunostaining of human thin (trunk) skin for Ki67, a common nuclear marker for cycling cells, and K15. Consistent with the data reported in Figure 1, K15 staining intensity is strongest in the bottom of rete ridges. The strongest Ki67 staining nuclei occur in keratinocytes showing low K15-intensity, and, frequently, a delaminating “fan-shaped” morphology (Fig. 2G) (see Miroshnikova et al., 2018).

Prior analyses of single-cell transcriptomics datasets established that *KRT15* is distinct from *KRT14* in the extent of co-regulation with a specific type II keratin partner. As reported (Cohen et al., 2022), transcription of *KRT14* and *KRT5* (R^2^=0.79), and similarly of *KRT10* and *KRT1* (R^2^=0.84), are tightly correlated over a broad range of expression levels (Fig. 2H). By contrast, *KRT15* shows a markedly weaker correlation to *KRT5* (R^2^=0.31; Fig. 2H) or any other type II keratin gene (data not shown). How the apparent imbalance in type I (*KRT14*, *KRT15*) and type II (*KRT5*) keratin transcripts is resolved at the protein level in progenitor keratinocytes of thin epidermis is an issue of interest but was not addressed in the current study.

Overall, the computational analyses reported here demonstrate that *KRT15* is highly expressed by cells with the strongest basal and weakest differentiated identity while *KRT10* is highly expressed by cells with the weakest basal and strongest differentiated identity. Further substantiating the distinct character of *KRT15*, the top 24 genes showing highest correlation to *KRT15* in human trunk skin (all cells) are highly enriched for components of hemidesmosomes and extracellular matrix (ECM) (Fig. 3A,B). In contrast, the top 24 genes most correlated to *KRT10* in human trunk skin (all cells) are highly enriched for desmosome components and proteins involved in terminal differentiation (Fig. 3A,C). These analyses confirm that *KRT15* and *KRT10* respectively define the progenitor vs. differentiating status while *KRT14* exhibits a broader distribution that spans both cellular states in the epidermis.

**Figure 3.**
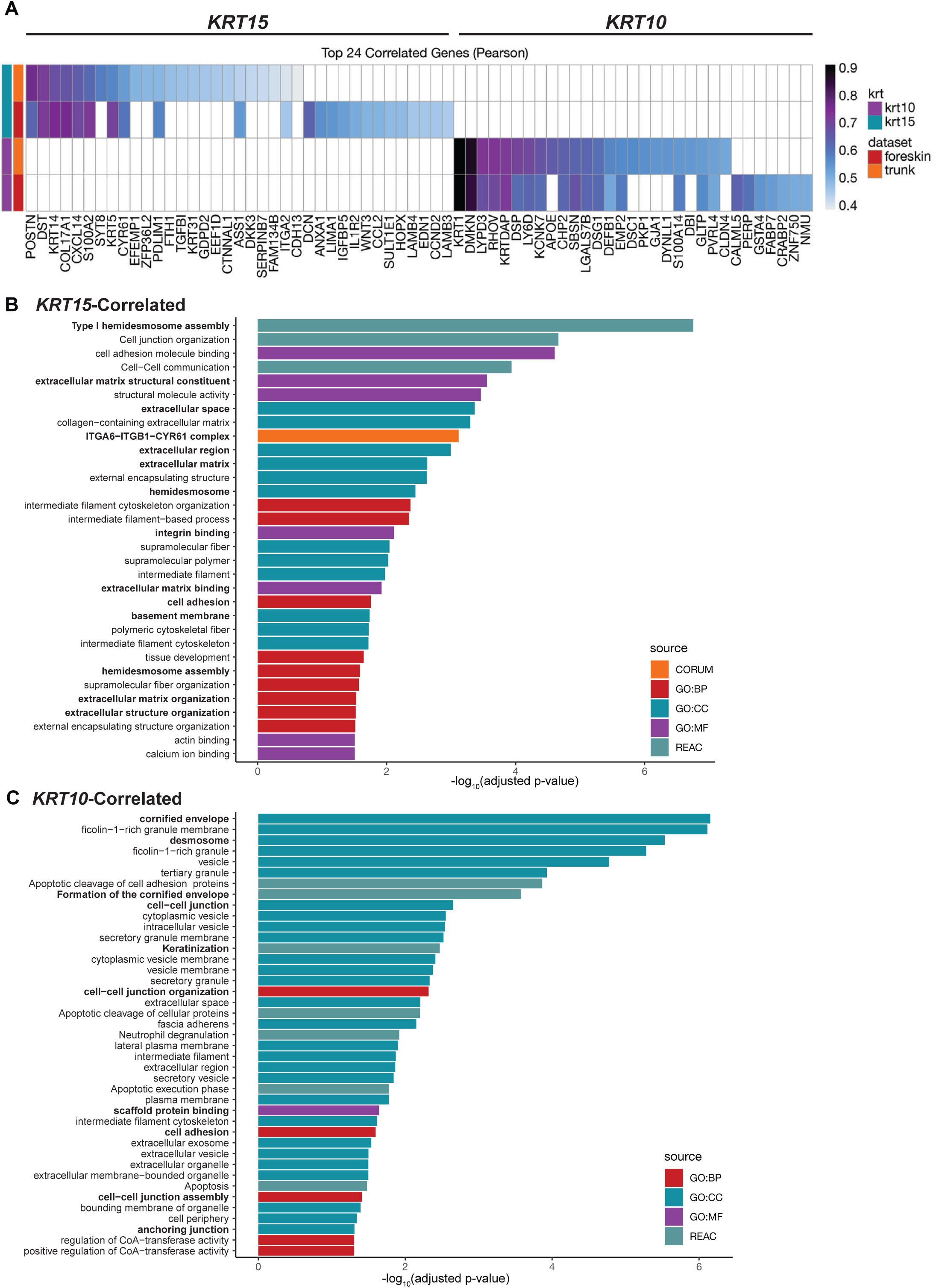
*KRT15* and *KRT10* expression define the extremities of the basal keratinocyte differentiation spectrum. **A.** Top 24 correlated genes for *KRT15* and *KRT10* in cells isolated from human trunk skin. **B.** Bar graph showing all GO categories enriched for *KRT15*-correlated genes with a cutoff of P < 0.05 from GProfiler. **C.** Bar graph showing all GO categories enriched for *KRT10*-correlated genes with a cutoff of P < 0.05 from GProfiler.

### Attributes of KRT15-expressing keratinocytes are also manifested in human foreskin and mouse back skin

Relative to trunk skin, *KRT15* is more prominently expressed in human foreskin (Suppl. Fig. 2A-D). Still, the basic attributes defining *KRT15* expression in human trunk (thin) skin (Fig. 2) apply to this tissue (Suppl. Fig. 2). Partitioning keratinocytes into quartiles according to the composite basal score (see Methods) highlights an enrichment for *KRT15* in keratinocytes within the quartiles showing the highest basal score and an enrichment for *KRT10* the quartiles showing the lowest basal score (Suppl. Fig. 2E). *KRT14*, on the other hand, manifests a hybrid character in this analysis (Suppl. Fig. 2E). A similar outcome is seen when organizing clusters according to the ratio of basal-to-differentiating scores (compare Suppl. Fig. 2F with Fig. 2F). Besides, an enrichment for G2-M^high^ keratinocytes in *KRT14^high^* keratinocytes is also maintained while a fraction of *KRT15^high^* keratinocytes show a high G2-M score in foreskin (Suppl. Fig. 2G). Further, relative to trunk skin, the correlation between *KRT5* and *KRT15* expression is stronger in foreskin (R^2^=0.43 vs. R^2^=0.31, respectively), while the *KRT5*-*KRT14* and *KRT1*-*KRT10* pairings still manifest a remarkably high correlation (R^2^=0.73 vs. R^2^=0.84, respectively; Suppl. Fig. 2 H,H’). While showing more prominent expression in foreskin, therefore, *KRT15* retains its character as a transcript that is enriched in a subpopulation of progenitor keratinocytes showing a strong basal character (Suppl. Fig. 2).

Similar analyses performed in a single cell transcriptomics data set obtained from mouse back skin (see Methods) show that the attributes that define *KRT15* expression in human trunk skin and foreskin are largely maintained in mouse skin compartment (Suppl. Fig. 3A-H’).

### A key role for the stutter cysteine in promoting the keratin-dependent sequestration of YAP1 to the cytoplasm

Previously we utilized transfection-permissive HeLa cells to show that K14 mediates the cytoplasmic retention of YAP1 through its stutter cysteine (Guo et al., 2020). To further assess the sufficiency of the stutter cysteine in conferring this property, we generated two variants, K15-A351C and K15 CF A351C, in which a cysteine was introduced in the second position of K15’s stutter motif. In the K15 CF A351C variant, all the cysteines occurring naturally in WT K15 have been mutated to alanines (designated “CF” for cysteine-free).

Consistent with our previous findings (Guo et al., 2020), co-expression of mCherry-K5 WT and EGFP-K14 WT (both human) in HeLa cells enabled filament polymerization and also resulted in localization of YAP1 to the cytoplasm and a decreased YAP1 nuclear:cytoplasmic ratio (Fig. 4A; quantitation reported in Fig. 4B). By contrast, co-expression of mCherry-K5 (human), EGFP-K14 WT (human), and (untagged, mouse) K15 WT reduced the ability of K14 WT to redistribute YAP1 to the cytoplasm (Fig. 4A,B). To test the hypothesis that K15’s ability to reduce the K14-dependent cytoplasmic sequestration of YAP1 is dependent on its lack of a stutter cysteine, we co-expressed mCherry-K5 and EGFP-K14 with either the K15 A351C or K15 CF A351C variant. Both K15 A351C and K15 CF A351C functioned as well as K14 WT in promoting YAP1 cytoplasmic sequestration (Fig. 4A,B), establishing that the stutter cysteine indeed plays a key role in mediating the cytoplasmic sequestration of YAP1.

**Figure 4.**
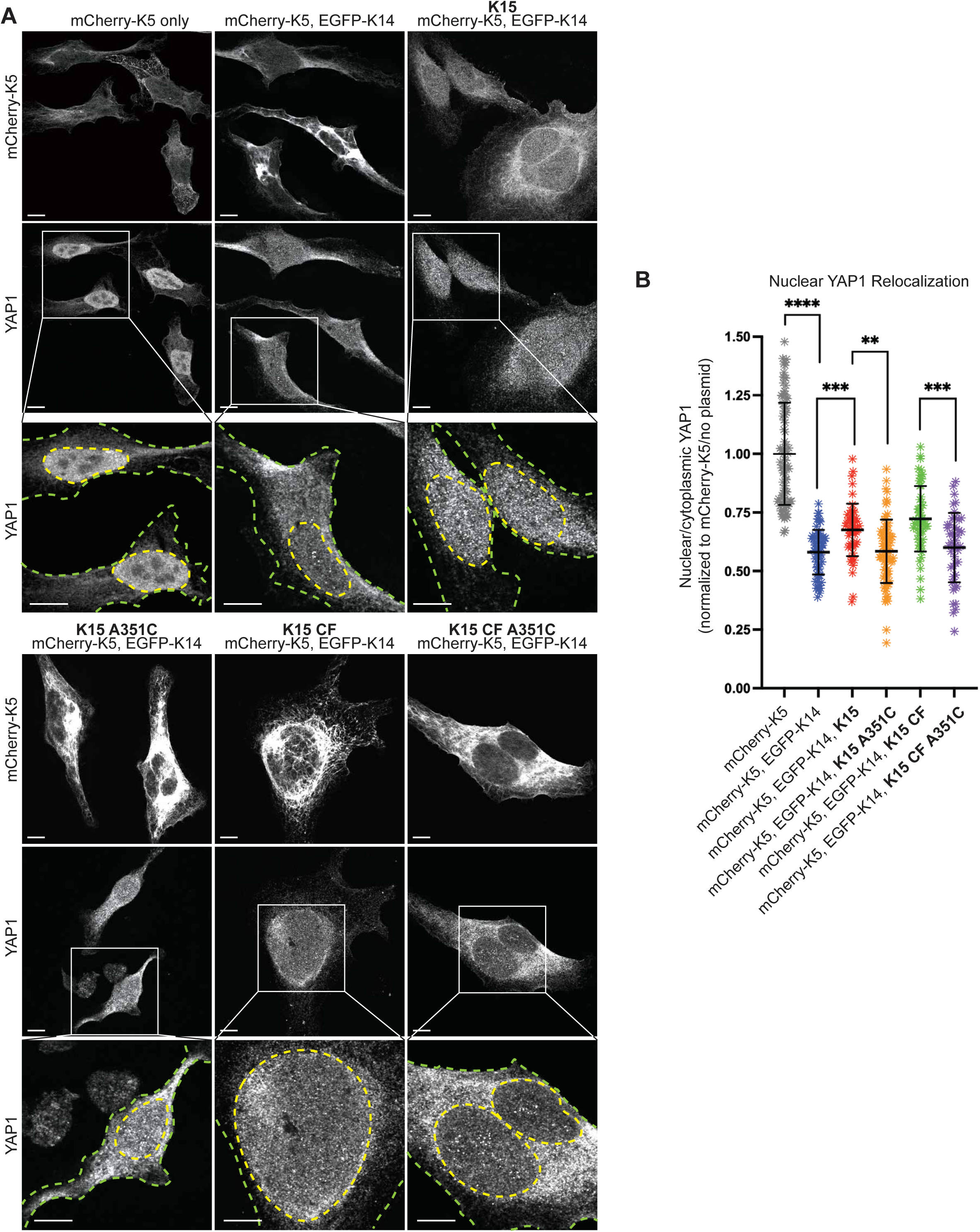
The stutter cysteine is sufficient to relocalize YAP1 in transfected HeLa cells. **A.** Representative images and magnified insets of HeLa cells transfected with mCherry-K5, EGFP-K14, and untagged WT and mutant K15. mCherry signal is autofluorescence, YAP1 is visualized through indirect immunostaining. Yellow dashed lines outline nuclei, green dashed line outline the cell peripheries. Scale bar, 10 µm. **C.** Scatter plots representing 3 pooled independent replicates, displaying mean and standard deviation. Dots represent a single cell. Comparisons made using Mann-Whitney tests. **** P <0.0001, *** P = 0.0002 and 0.0003, ** P = 0.0083.

We next sought to test the impact of K15 expression on K14-dependent relocalization of YAP1 to the cytoplasm of mouse skin keratinocyte cultures. Skin keratinocytes obtained from newborn *Krt14^WT/null^* hemizygous mouse pups were seeded for primary culture and transfected with EGFP-tagged (human) K14, K15, or K15 A351C overexpression vectors. Transfected cells were expanded for 2 days in calcium-free basal growth media before being switched to growth media containing 1.2 mM CaCl_2_, a physiological trigger of differentiation. Cells were calcium-treated for 24 hours, fixed, and stained for YAP1. Keratinocytes in primary culture display tight cell-cell adhesions and overlapping segments of cytoplasm. Resolving cytoplasmic YAP1 signal in individual cells is therefore unfeasible, and we instead opted to specifically quantitate nuclear YAP1 as the key metric of interest. As expected, nuclear YAP1 immunostaining decreased in mock (expressing endogenous K14 WT and K15 WT) and EGFP-K14 transfected keratinocytes following calcium-induced differentiation (Fig. 5A, with magnified insets of nuclear YAP1 shown in Fig. 5A’; quantitation shown in Fig. 5B).

**Figure 5.**
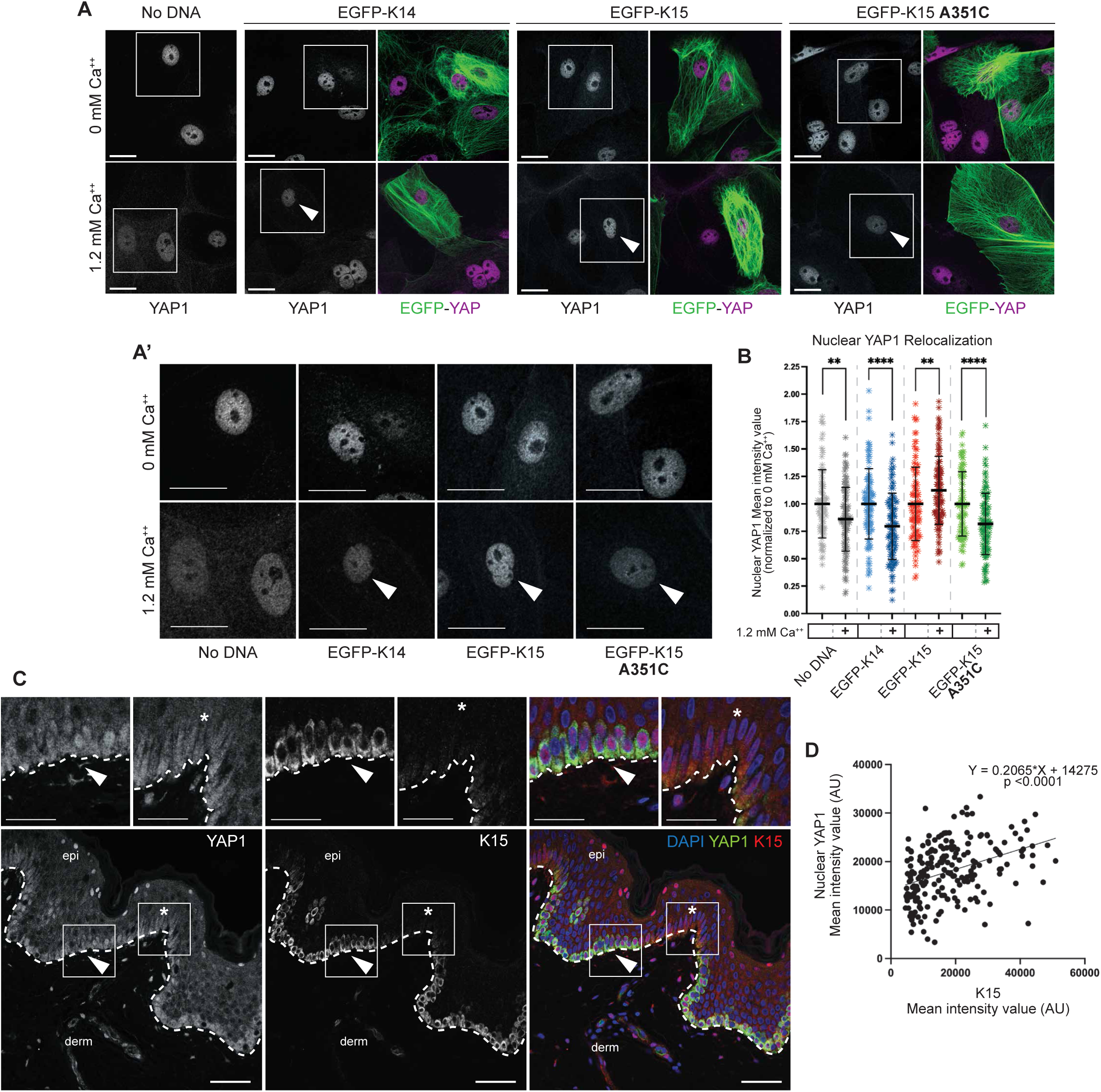
The stutter cysteine is sufficient to relocalize YAP1 in differentiating mouse skin keratinocytes. **A.** Representative micrographs of *Krt14^WT/null^* keratinocytes in primary culture transfected with EGFP-K14, EGFP-K15, or EGFP-K15 A351C, with and without 1.2 mM CaCl_2_ to induce differentiation. YAP1 is visualized through immunofluorescence. “Merge” panels display autofluorescent EGFP (green) and YAP1 (magenta). Arrowheads indicate EGFP-positive keratinocytes. Scale bar, 10 µm. **A’.** Magnified insets of nuclei with YAP1 indirect immunofluorescence from Figure 5A. Arrowheads indicate EGFP-positive keratinocytes. Scale bar, 10 µm. **B.** Scatter plots representing 3 independent replicates, pooled, displaying mean ± SD. Dots represent a single cell. Comparisons made using unpaired t tests with Welch’s correction. **** P <0.0001, ** P = 0.0058 and 0.0014. **C.** Multiplexed immuno-staining for K15 (green) and YAP1 (red). Nuclei are counterstained with DAPI (blue). Dashed line specifies the basal lamina. epi, epidermis; derm, dermis. Scale bar, 50 µm. Arrowheads depict region with strong K15 and nuclear YAP1 staining. **D.** XY scatter plot is comparing nuclear YAP1 intensity (arbitrary units) vs. K15 intensity (arbitrary units). Each dot represents a single basal keratinocyte. Equation for the simple linear regression of the comparison is displayed, showing a positive correlation. Regression is significantly nonzero (P <0.0001).

By contrast, transfection of EGFP-K15 antagonized the calcium-induced reduction of nuclear YAP1 signal (Figs. 5A,A’,B), in agreement with the HeLa, *in vivo* morphological, and computational data presented above. On the other hand, overexpression of knock-in stutter cysteine EGFP-K15 A351C resulted in loss of nuclear YAP1 signal in response to calcium as it did in mock and EGFP-K14 WT transfected cells (Figs. 5A,A’,B). The outcome of transfection assays involving human HeLa cells and mouse keratinocytes in primary culture prompted the prediction that basal layer keratinocytes expressing K15 at higher levels should also exhibit stronger partitioning of YAP1 to the nucleus in the human epidermis *in situ*. Dual staining for K15 and YAP1 in sections of thin human skin epidermis reveals a significant and positive correlation between K15 signal intensity and nuclear YAP1 signal intensity (R^2^=0.15, P<0.0001; Fig. 5C, quantitation in Fig. 5D). Further, nuclear YAP1 signal intensity is, similarly to K15, strongest at the bottom of rete ridges (Fig. 5C). Together this in vitro, ex vivo, and in vivo data provides direct evidence that K15, through its lack of a stutter cysteine, is poised to maintain a progenitor cellular state by promoting nuclear YAP1 localization.

### A candidate 14-3-3 binding site in the N-terminal head domain of K14 is missing in K15

We next sought to assess whether K15 and K14 differ in their ability to sequester YAP1 to the cytoplasm in relation to the property of 14-3-3 binding. Previous studies have shown that the 14-3-3σ isoform, in particular, promotes in the sequestration of YAP1 in the cytoplasm at an early stage of keratinocyte differentiation in interfollicular epidermis (Sambandam et al., 2015; Sun et al., 2015). Moreover, co-immunoprecipitation and PLA assays have shown that K14 physically associates with, and is spatially proximal to, 14-3-3σ in newborn mouse skin keratinocytes (Guo et al., 2020), though the cis-acting determinant(s) involved remain unknown. On the other hand, using proximity ligation assays, Ievlev et al. (2023) reported that, unlike K14, K15 lacks spatial proximity to 14-3-3σ in human surface airway epithelial cells in culture and concluded that K15 does not bind 14-3-3σ and thus cannot regulate YAP1 like K14 does.

14-3-3 adaptor proteins typically bind their client proteins, including YAP1 (Pocaterra et al., 2020), in a phosphorylation-dependent manner (Pennington et al., 2018). Many IF proteins, in addition to K14, can bind 14-3-3 adaptors. For human K17 (Kim et al., 2006), human K18 (Ku, 1998), *Xenopus* K19 (Mariani et al., 2020) and human vimentin (Tzivion et al., 2000), studies have shown that specific serine (S) residues located in the N-terminal head domain mediate the interaction with 14-3-3 adaptors. The *14-3-3-Pred* software (Madeira, 2015) predicts two strong potential binding sites in human K14, S33 and S44, with S44 receiving the highest score (Suppl. Fig. 4A). S44 in human K14 is orthologous to S44 in K17, previously shown to mediate 14-3-3σ binding (Kim et al., 2006). Analyzing the mouse K14 sequence via *14-3-3-Pred* identifies S40 (Fig. 1A), which is orthologous to S44 in human K14, as the most probable 14-3-3 binding site (Suppl. Fig. 4A,A’). Finally, our mass spectrometry analyses show that K14’s S39 and S44 are phosphorylated in human N-TERT keratinocytes in culture (Suppl. Fig. 4D,D’). Given this evidence, we hypothesized that the S^39^CRAPS^44^ motif in human K14’s N-terminal head domain (see Fig. 1A), which is perfectly conserved in mouse, mediates interaction with 14-3-3α. The corresponding sequence in human and mouse K15, G^29^FGGGS^34^ (numbering for the human ortholog; see Fig 1A), is markedly different. Accordingly, we generated a human K14 variant in which the SCRAPS motif in the N-terminal head domain is mutated to GFGGGS, and a human K15 variant in which GFGGGS is replaced with K14’s SCRAPS motif. These two newly generated variants were tested for their ability to regulate the subcellular distribution of YAP1 and their spatial proximity and physical interaction with 14-3-3 in epithelial cells in culture.

HeLa cells were co-transfected with K5 and either EGFP-K14 WT, EGFP-K15 WT or EGFP-K15-SCRAPS-A351C and initially processed for immunoprecipitation of GFP, a shared determinant, from the detergent-soluble pool (see Methods). 14-3-3σ was reproducibly detected in GFP immunoprecipitates from all three transfection combinations, indicating that it physically interacts with K14, K15, and K15 SCRAPS A351C (Fig. 6A; quantitation shown in Fig. 6B). However, the amount of EGFP-K15 WT protein detected in the soluble pool as well as in the GFP immunoprecipitate fractions was significantly larger than that detected in EGFP-K14 WT and EGFP-K15 SCRAPS A351C proteins (Fig. 6A,B), suggesting that WT K15 protein is more readily detergent soluble. To address this bias, we normalized the signal for 14-3-3σ against that of GFP and the resulting ratios suggest that there was >2-fold more 14-3-3σ in the EGFP-K14 WT and EGFP-K15 SCRAPS A351C immunoprecipitates than in the EGFP-K15 WT immunopecipitate (see “14-3-3σ enrichment” chart in Fig. 6B). Such findings show that, consistent with the 14-3-3Pred analysis reported in Suppl. Fig. 4A,A’,C,C’, both K14 and K15 can physically associate with 14-3-3σ, directly or indirectly. When factoring in the differences observed in IP detergent solubility, they also suggest that K15 has a lower affinity for the relevant 14-3-3σ-containing complexes that is remediated by replacing “GFGGGS” with “SCRAPS” in its head domain. Finally, we cannot exclude that K5 plays a role in the interaction with 14-3-3σ in this setting – of note, two specific sites in each of human and mouse K5 are predicted to confer 14-3-3 binding (Suppl. Fig. 4B,B’).

**Figure 6.**
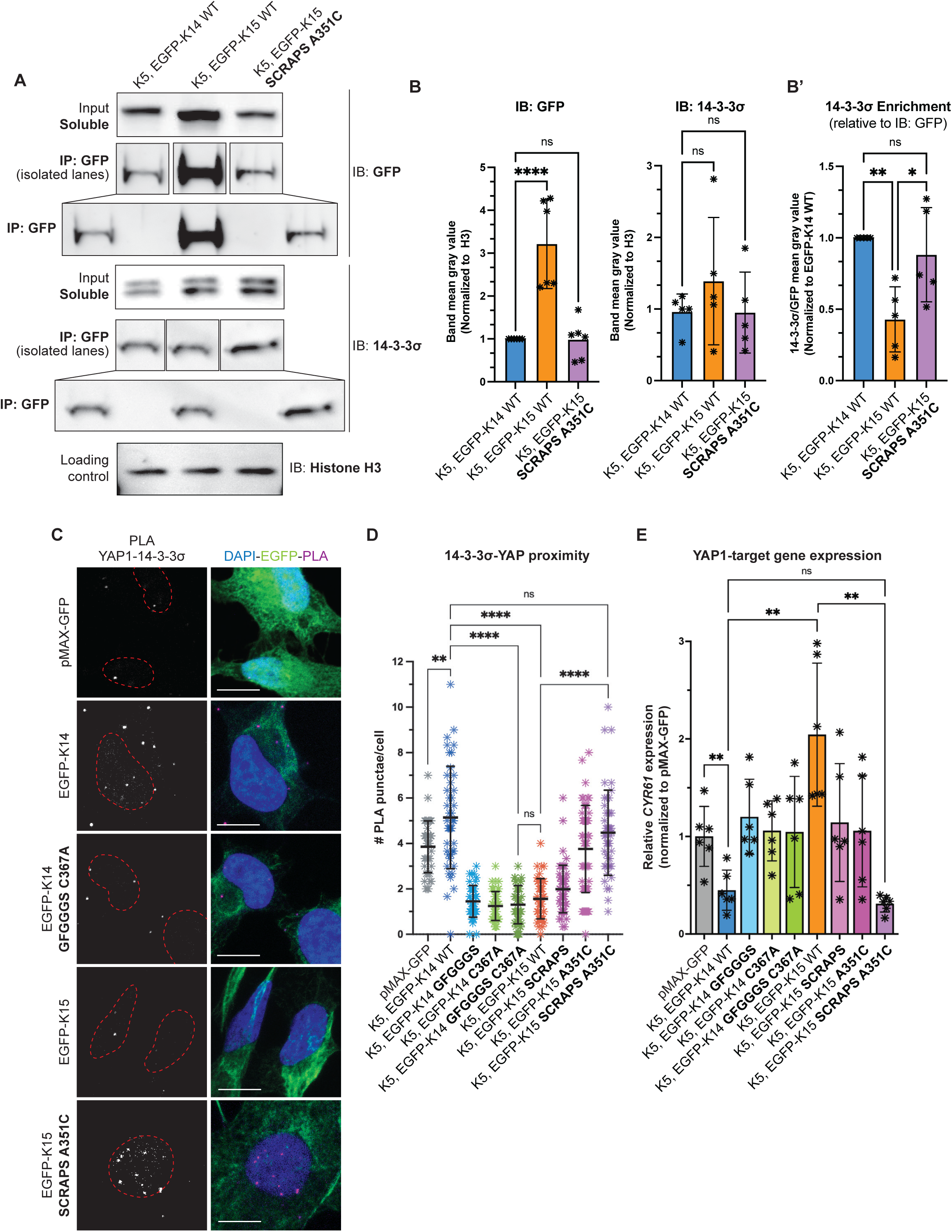
The SCRAPS motif and the stutter cysteine are required to interconvert YAP1-interaction between K14 and K15. **A.** Representative co-immunoprecipitations of HeLa cells transfected with untagged K5 and WT K14, K15, and mutant K15. Pulldown of endogenous 14-3-3σ is depicted. **B.** Differential levels of WT K15, relative to WT K14 or mutant K15, were observed in the soluble fraction. Transfection of keratin(s) did not affect endogenous 14-3-3σ levels. WB intensity, normalized to loading control (Histone H3). Dots represent 3 independent replicates, displaying mean and standard error. **B’.** Endogenous 14-3-3σ pulls down more efficiently with WT K14 and mutagenized K15, as compared to WT K15. Equation to calculate 14-3-3σ pulldown efficiency is described in Methods. Dots represent 3 independent replicates. **C.** Representative micrographs of YAP1-14-3-3σ proximity ligation assay performed on transfected HeLa cells. Cells were transfected with untagged keratin 5 and WT and mutant K14 or K15. Merge panels show PLA punctae (red), EGFP-tagged keratin autofluorescence (green), and DAPI counterstain (blue). Scale bar = 10 µm. **D.** Scatter plots representing YAP1-14-3-3σ PLA punctae counted per cell in 3 pooled independent replicates, displaying mean and standard deviation. Dots represent a single cell. Comparisons made using Mann-Whitney tests. **** P <0.0001, ** P = 0.0033, ns = not significant. **E.** Transcription of a YAP1 target gene, *CYR61*, as measured via RT-qPCR in HeLa cells transfected with untagged keratin 5 and WT and mutant K14 or K15. Dots represent 6 independent replicates. Comparisons made using Mann-Whitney tests. **** P <0.0001, ** P = 0.0033, ns = not significant.

We next compared the ability of K14, K15, and their variants to promote spatial proximity between YAP1 and 14-3-3σ in transfected HeLa cells using proximity ligation assays (PLA). Co-expression of untagged K5 WT (human) and EGFP-K14 WT (human) in HeLa cells resulted in a robust PLA signal when probing for YAP1 and 14-3-3σ (Fig. 6C; signal quantitation reported in Fig. 6D). By comparison, co-expression of mCherry-K5 and either EGFP-K14-C367A, EGFP-K14-GFGGGS, or EGFP-K14-GFGGGS-C367A double mutant was ineffective at promoting YAP1/14-3-3σ physical proximity (Fig. 6C,D). These findings confirm previous studies of the K14 C367A variant (Guo et al., 2020) and support the notion that the SCRAPS motif in K14’s N-terminal head plays a role in the subcellular partitioning of YAP1 via 14-3-3σ interaction. By contrast, relative to EGFP-K14 WT, co-expression of K5 WT and EGFP-K15 WT is as ineffective as EGFP-K14-C367A, EGFP-K14-GFGGGS, and EGFP-K14-GFGGGS-C367A at promoting YAP1/14-3-3σ physical proximity (Fig. 6C,D). Addition of both the SCRAPS motif and the stutter cysteine to K15 (EGFP-K15-SCRAPS-A351C double-mutant) yielded a YAP1/14-3-3σ PLA signal that is statistically the same as EGFP-K14 WT (Fig. 6C,D). While EGFP-K15-A351C was able to promote an intermediate level of YAP1/14-3-3σ proximity, the EGFP-K15-SCRAPS variant was equivalent to EGFP-K15 WT (Fig. 6C,D). Such PLA findings suggest that the stutter cysteine and the SCRAPS motif each contribute significantly to promoting spatial proximity between K14 and YAP1, mediated via 14-3-3σ, in the cytoplasm, while the capacity of WT K15 to do so is markedly impaired because it lacks these two specific determinants.

Finally, to examine whether these cytoplasmic interaction(s) impair YAP1 transcriptional activity, we assessed the transcriptional output of YAP1 by measuring *CYR61* mRNA levels using RT-qPCR in transfected HeLa cells. *CYR61*, also known as *CCN1*, is a bona fide and robust YAP1 target gene; see (Ma et al., 2017). The reference in this assay is *CYR61* measured in cells transfected with pMAX-GFP alone (set at a value of 1 across replicates; see Fig. 6E). As previously shown (Guo et al., 2020), cells co-transfected with K5 and EGFP-K14WT show a dramatic reduction in *CYR61* transcripts, while cells co-transfected with K5 and EGFP-K14 C367A do not (Fig. 6E), setting up the stage for assessing the properties of variants in K14 and K15 WT. We found that EGFP-K14 GFGGGS and EGFP-K14 GFGGGS C367A were equally ineffective at attenuating the transcription of *CYR61* (Fig. 6E and Supp. Fig. 4E), showing that replacing “SCRAPS” with “GFGGGS” in the head domain of K14 abrogates the latter’s intrinsic ability to mitigate YAP1-dependent transcription. Remarkably, *CYR61* transcripts were markedly higher in cells co-transfected with K5 and EGFP-K15 WT compared to the pMAX-GFP reference (Fig. 6E), indicating that K15 is unable to negatively regulate YAP1 (a finding that is consistent with the PLA data reported in Fig. 6D). While each of the EGFP-K15 SCRAPS and EGFP-K15 A351C variants performed better than EGFP-K15 WT in this assay (Supp. Fig. 4E), only EGFP-K15 SCRAPS A351C was as effective as EGFP-K14 WT in attenuating the transcriptional activity of YAP1 when co-transfected with K5 in HeLa cells (Fig. 6E). These findings confirm and extend the functional relevance of the PLA data (see Fig. 6D) in clearly demonstrating that K14 and K15 markedly differ in their ability to control YAP1 activity, correlating tightly with regulation of its subcellular localization and spatial proximity to 14-3-3σ. Mechanistically, these findings confirm a key role for the stutter Cys in coil 2 of the rod domain and uncover a role for the SCRAPS motif in N-terminal head domain of K14 for the property of YAP1 regulation. This said, our findings also highlight the complexity of the interaction between keratins and 14-3-3σ with regards to this role (see Discussion). Finally, we cannot formally rule out that the manipulation of K14 and K15 sequences using mutagenesis, along with the addition of GFP tags, have had unintended consequences that contributed to the outcome of our studies.

### K15 overrepresentation alters basal keratinocytes and the basement membrane zone in the epidermis

The experimental and computational data reported so far substantiate the notion that keratinocytes with higher levels of K15, relative to K14, possess a stronger progenitor identity. To test this hypothesis *in vivo* we sought to manipulate the relative levels of K15 and K14 proteins in mouse skin. To do so we generated a mouse strain harboring compound heterozygous alleles at the *Krt14* locus - *Krt14^C373A/null^*. This strain possesses a single allele of *Krt14* that is mutated at the stutter cysteine, while the *Krt15* alleles are unperturbed.

*Krt14^C373A/null^* mice are viable and blister-free (see below), indicating they express sufficient K14 to maintain epithelial integrity. To assess relative levels of K14 to K15, we performed quantitative dot blotting on whole tail skin lysate harvested from 8 week-old male *Krt14^C373A/null^* mice, with their *Krt14^C373A/WT^*littermates used as control. *Krt14^C373A/null^* tail lysate displayed a significant 25.3% reduction in K14 mean steady state levels (Suppl. Fig 5A). K15, in contrast, was not significantly different between *Krt14^C373A/WT^* and *Krt14^C373A/null^* samples (Suppl. Fig 4A’), confirming the alteration to the K14 to K15 ratio in *Krt14^C373A/null^* skin.

In the progeny resulting from crosses between *Krt14 ^C373A/C373A^* and *Krt14^WT/null^* mice, we noticed that a subset of pups, by P4, had a plumper, darker and slightly larger appearance as the first group of hair follicles completes differentiation and fur emerges at the skin surface (Fig. 7A). Upon weighing, such pups were on average 10% heavier than their littermates (3.02g vs. 2.75g; Suppl. Fig.5B). Genotyping performed at weaning indicated that these features were completely specific to the *Krt14^C373A/null^* genotype, with no sex bias. The difference in body weight does not persist after pups were weaned, such that *Krt14^C373A/null^* and *Krt14^C373A/WT^* mice were indistinguishable as young adult mice.

**Figure 7.**
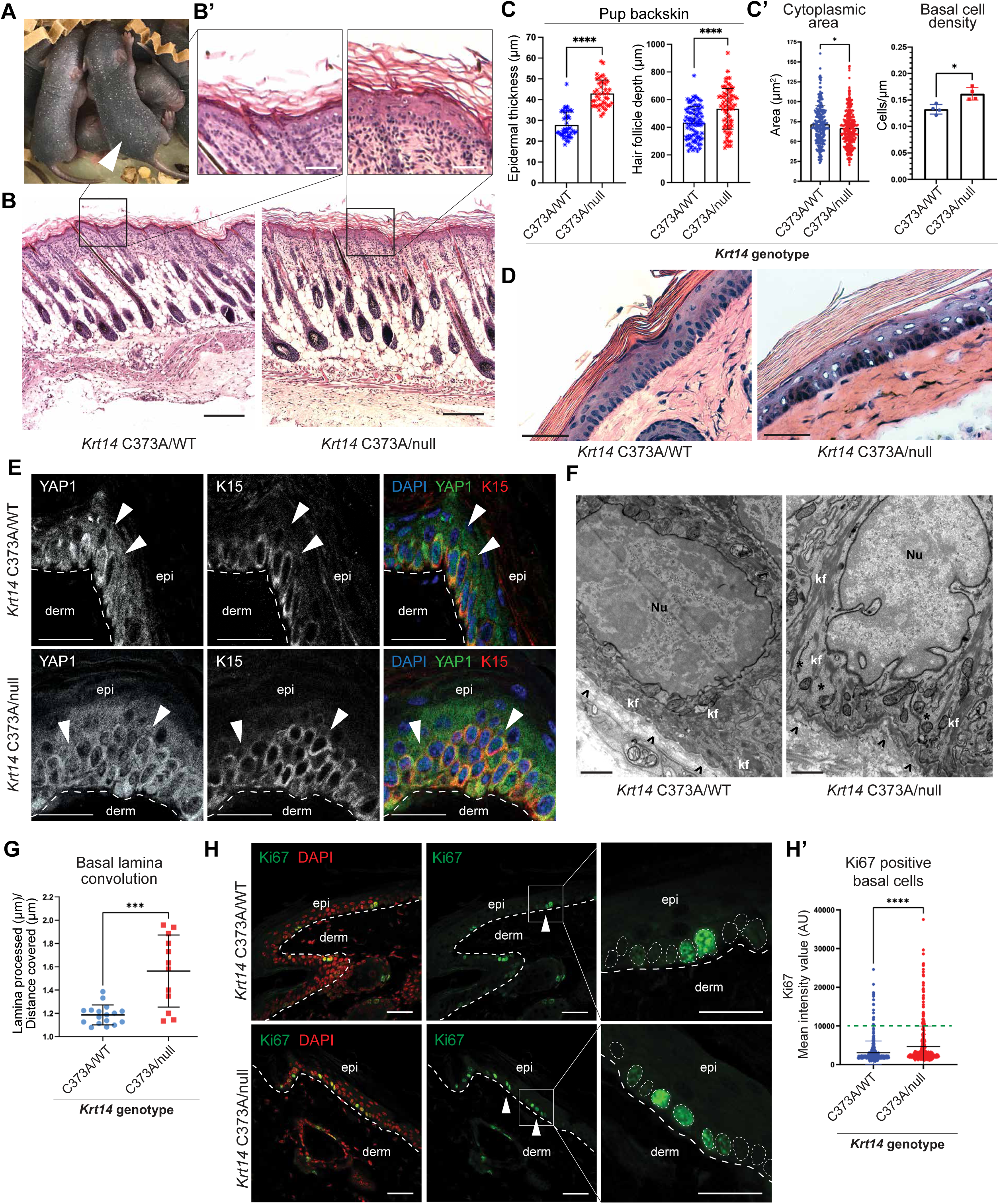
K15 overrepresentation alters defining attributes of basal layer keratinocytes *in vivo*. **A.** *Krt14^C373A/WT^* and *Krt14^C373A/null^* (arrow) littermates at P5. **B.** H&E staining performed on back skin harvested from P5 *Krt14^C373A/WT^* and *Krt14^C373A/null^* littermate pups. Scale bar, 50 µm. **B’.** Magnified insets of H&E stained back skin in Figure 5A. Scale bar, 10 µm. **C.** Epidermal thickness (µm) and hair follicle depth (µm) in back skin harvested from P5 *Krt14^C373A/WT^*and *Krt14^C373A/null^* littermate pups. Comparisons made using Mann-Whitney tests. **** P <0.0001. **C’.** Mean cytoplasmic area of basal keratinocytes with standard deviation; dots represent single cells. Comparisons made using Mann-Whitney tests. * P = 0.0286. Basal cell density calculated by mean cells/µm of basal lamina with standard deviation; dots represent individual micrographs from tail skin harvested from 4 littermate animals each. Comparisons made using Mann-Whitney tests. * P = 0.014. **D.** H&E staining performed on tail tissue harvested from 8 week old *Krt14^C373A/WT^*and *Krt14^C373A/null^* tail tissue. Scale bar = 50 µm. **E.** YAP1 and K15 immunofluorescence performed on 8 week old *Krt14^C373A/WT^*and *Krt14^C373A/null^* littermate tail skin. Merge panel displays DAPI counterstain (blue), YAP1 (green), and K15 (red). Arrows emphasize nuclei of suprabasal keratinocytes, showing YAP1-negative nuclei in *Krt14^C373A/WT^* tail skin and YAP1-positive nuclei in *Krt14^C373A/nul^* tail skin. Dashed line specifies the basal lamina. epi, epidermis; derm, dermis. Scale bar, 50 µm. **F.** Representative transmission electron microscopy micrographs of 8 week old *Krt14^C373A/WT^* and *Krt14^C373A/null^* basal keratinocytes in ear skin. Nu, nucleus; kf, keratin filaments; arrows, hemidesmosomes; *, areas devoid of keratin filaments. Scale bar, 800 nm. **F.** Ki67 immunofluorescence performed on 8-week old *Krt14^C373A/WT^* and *Krt14^C373A/null^*littermate tail skin. Merge panel displays Ki67 (green) with DAPI counterstain (red). Arrows indicate Ki67-positive basal nuclei. Dashed line specifies the basal lamina. epi, epidermis; derm, dermis. Scale bar, 50 µm. **G.** Mean of the basal lamina processed/ distance covered to represent basal lamina convolution, displayed with standard deviation. Dots are individual micrograph images. Comparisons made using Mann-Whitney tests. *** P = 0.0009. **G’.** Mean of the Ki67 mean intensity value for individual basal keratinocyte nuclei, displayed with standard deviation. Percentages of cells above the visual threshold for Ki67 positivity (green dashed line). Comparisons made using Mann-Whitney tests. **** P <0.0001.

The plumper appearance of the *Krt14^C373A/null^* pups suggested that they may exhibit thicker skin, a suspicion that was confirmed through routine histology (Fig. 7B, magnified inset in 7B’). The increase in *Krt14^C373A/null^* skin thickness was most significant in the dermis and correlated with a general increase in hair follicle length (by 19%, on average; Fig. 7C). The latter is consistent with the observation, reported above, that hair erupts a day earlier in *Krt14^C373A/null^* relative to *Krt14^C373A/WT^*littermates. Quantitation showed that the interfollicular epidermis was also thickened, by 35%, in the *Krt14^C373A/null^* pups (Fig. 7C).

Upon closer examination, additional alterations were observed in the epidermis of hematoxylin-eosin-stained sections of *Krt14^C373A/null^*skin at 8 weeks. In the upper epidermis of the tail, the granular layer was reduced in both density and contrast while the cornified layers were expanded, respectively suggesting altered terminal differentiation and hyperkeratosis. In the lower epidermis, basal keratinocytes were crowded and showed clustering in *Krt14^C373A/null^*mice whereas they were evenly distributed along the basal lamina in *Krt14^C373A/WT^*controls. Areas with nuclear clustering displayed muddled H&E staining (Fig. 7D). To quantitate these features, we measured the mean surface area of basal keratinocytes and their density per unit length of dermo-epidermal interface. The mean area of *Krt14^C373A/null^* basal keratinocytes was significantly smaller than *Krt14^C373A/WT^* basal keratinocytes in tail epidermis (*Krt14^C373A/WT^*: 71.50 µm^2^ vs. *Krt14^C373A/null^*: 66.70 µm^2^; Fig. 7C’). In concurrence, the density of basal keratinocytes per µm of dermo-epidermal interface was significantly increased in *Krt14^C373A/null^*keratinocytes (*Krt14^C373A/WT^*: 0.1324 cells/µm vs. *Krt14^C373A/null^*: 0.1616 cells/µm, Fig. 7C’).

Consistent with previous reports on homozygous *Krt14^C373A/C373A^* mouse tail skin (Guo et al., 2020), the tail skin epidermis of young adult *Krt14^C373A/null^* mouse displays aberrant suprabasal nuclear YAP1 staining, in contrast to *Krt14^C373A/WT^*epidermis in which nuclear YAP1 staining is restricted to the basal layer (Fig. 7E). Moreover, K15 staining atypically extended into the suprabasal layers of *Krt14^C373A/null^*tail skin (Fig. 7E). As expected and in concurrence with previous reports (Guo et al., 2020), transcription of the YAP1-target gene *CYR61* is also significantly increased in the tail skin epidermis of young adult *Krt14^C373A/null^*compared to *Krt14^C373A/WT^* littermates (Supp. Fig. 5C).

We next used transmission electron microscopy (TEM) to identify any ultrastructural alterations in *Krt14^C373A/null^* epidermis. We imaged skin dissected from the back, ear, and tails of 8 week-old *Krt14^C373A/null^*and *Krt14^C373A/WT^* littermates, male and female (Fig. 7F). Recurring observations from *Krt14^C373A/null^* skin include alterations to nuclear morphology and an increased incidence of cytoplasmic regions devoid of keratin filaments. The basal lamina in *Krt14^C373A/null^* basal keratinocytes also displays an increase in its spatial convolution and its thickness (Fig. 7G). Additional recurrent features specific to *Krt14^C373A/null^*basal keratinocytes include more prominent hemidesmosomes, typically associated with distortions in the basal lamina, along with cytoplasmic regions devoid of keratin filaments (Fig. 7F; see “kf”). These analyses show that K15 overrepresentation is associated with alterations to the morphological attributes of basal keratinocytes, hemidesmosomes and the basal lamina, potentially influencing cell cycling and development, and providing *in vivo* evidence for a pro-progenitor role for K15 in epidermis. Whether the striking alterations observed in the skin of *Krt14^C373A/null^* mice involves the misregulation of other effectors, in addition to YAP1, is likely but unclear at this time.

Finally, to complement the data related in Figure 2G and test whether basal cell crowding in adult *Krt14^C373A/null^* epidermis is associated with increased cell divisions, we performed Ki67 immunofluorescence staining, a highly specific marker for cycling cells, on tail skin harvested from 8 weeks old *Krt14^C373A/WT^* and *Krt14^C373A/null^* male and female littermates. As expected, Ki67 staining is exclusively localized to the basal keratinocytes in both *Krt14^C373A/WT^* and *Krt14^C373A/null^* tissue. *Krt14^C373A/null^* tissue, however, showed a significant increase in Ki67 mean intensity value in basal nuclei (Fig. 7H). We next set a threshold for Ki67 positivity by determining the median Ki67 intensity for nuclei that were visually negative for Ki67 and calculated the percentage of basal cells above this threshold. In *Krt14^C373A/WT^* basal keratinocytes, 6.8% of nuclei were Ki67-positive, while 16.8% of *Krt14^C373A/null^*basal keratinocyte nuclei were Ki67-positive (Fig. 7H’). Both parameters confirm an increase in Ki67-positive nuclei in 8 weeks old *Krt14^C373A/null^* tail tissue compared to *Krt14^C373A/WT^*littermates.

## DISCUSSION

A link between K15 expression and an epithelial stem character was first uncovered in the human hair bulge by Lyle *et al*. more than 25 years ago (Lyle et al., 1998). This association has since been confirmed and expanded to mouse skin (Liu et al., 2003) as well as to internal stratified epithelia, e.g., the esophagus (Giroux et al., 2017) (reviewed in (Bose et al., 2013)). Follow-up studies, in human and in mouse, have shown that *Krt15^high^*-expressing keratinocytes residing in the basal layer (and purified based on high surface levels of integrins) express markers associated with stemness (e.g., CD34, CD200, integrin B1^bright^, Lgr5) (Inoue et al., 2009; Jaks et al., 2008; Morris et al., 2004; Trempus et al., 2003; Tumbar et al., 2004) and are capable of reconstituting all epithelial lineages in mature hair follicles (Morris et al., 2004) and esophagus (Giroux et al., 2017). In striking contrast, *Krt15^low^*-expressing, basally-located keratinocytes (which are otherwise *Krt14^high^* in character) do not exhibit this lineage potential (Morris et al., 2004). In human epidermis, K14 and K15 are co-expressed in keratinocytes at the transcript and protein levels, but the distribution of K15 is unique in many respects; unlike K14, K15 occurs in a subset of basal keratinocytes that are preferentially located in the deeper area of epidermal rete ridges, and does not persist in suprabasal keratinocytes (see Jonkman et al., 1996; Porter et al., 2000; Waseem et al., 1999; Webb et al., 2004; Whitbread and Powell, 1998; Zhan et al., 2007). We confirmed and significantly extended these observations via analyses of recent single cell transcriptomics data and targeted immunostainings. As intriguing as these attributes are, however, there is, as of yet, no biochemical or mechanistic insight that accounts for the intriguing connection between K15 expression and an epithelial stem character in skin.

Here, we report on findings showing that, owing to the lack of two key cis-acting determinants - a cysteine within the stutter of the α-helical central rod domain and the SCRAPS motif within the N-terminal head domain - K15 is unable to effectively mediate the cytoplasmic sequestration of YAP1. Accordingly, and unlike K14 (Guo et al., 2020), K15 may be unable to enact a switch in Hippo signaling, from “off” to “on”, prompting progenitor basal keratinocytes to initiate differentiation in the epidermis (Yuan et al., 2022; Zhang et al., 2011). **Figure 8** illustrates a model that integrates these new findings with our previous work focused on the K14/14-3-3σ/YAP1 interaction (Guo et al., 2020). The model proposes that a higher K15:K14 protein ratio in basal keratinocytes of the epidermis promotes the progenitor state and antagonizes their differentiation. Once *KRT15* expression subsides and the K15:14 protein ratio falls below a postulated threshold (a reality favored by K15’s apparent shorter half-life (Cui et al., 2022), its higher solubility (this study), along with persistence of *KRT14* expression at significant levels), K14 is able to effectively sequester YAP1 to the cytoplasm (after its specification through post-translation modifications) and help promote the initiation of keratinocyte differentiation.

**Figure 8.**
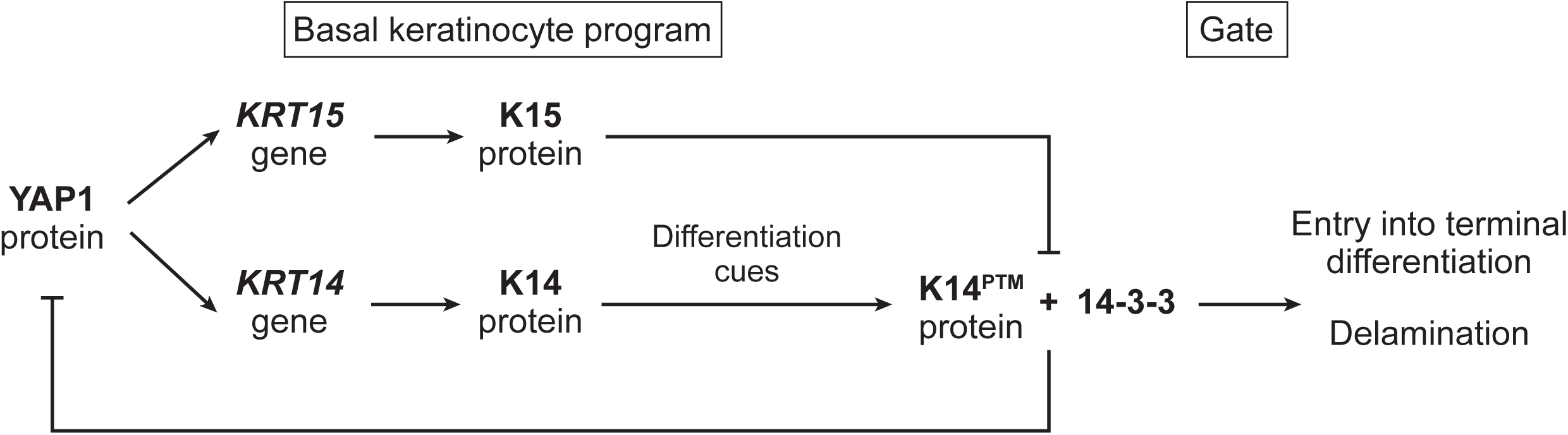
Model. Interplay between keratin 15 and keratin 14 in regulating YAP1’s access to the nucleus in progenitor keratinocytes of the epidermis. The model proposes that a higher K15:K14 protein ratio in basal keratinocytes of epidermis promotes the progenitor state and antagonizes differentiation. See text for details.

The success of our efforts to interconvert, through mutagenesis, the ability of K14 and K15 to regulate the subcellular partitioning and transcriptional role of YAP1 in keratinocytes highlights a dominant role for the stutter cysteine located in coil 2 of the central α-helical rod domain. We note that, in addition to K15, two additional type I keratins associated with a stem-like epithelial character in skin, namely K19 (Michel, 1996) and K24 (Wang et al., 2025), also lack the stutter cysteine, while the differentiation specific K10 and K9 (glabrous skin) feature it (Guo et al., 2020; Lee et al., 2012). A recent effort in our laboratory focused on glabrous skin showed that, as we predicted in Guo et al. (2020), K9 and its stutter cysteine are required for the cytoplasmic sequestration of YAP1 in differentiating keratinocytes of glabrous skin in vivo and epithelial cell culture ex vivo (Steiner et al., 2025). The identity of the K14 stutter cysteine-dependent disulfides, and the mechanism(s) of their formation, have emerged as open issues of significant interest(see Guo et al., 2020, for discussion).

Besides extending the crucial role of the stutter cysteine in coil 2 of the rod domain, the findings we report here also point to a novel role for the SCRAPS motif in the N-terminal head domain of K14 towards regulating the subcellular partitioning and role of YAP1. We also show that the SCRAPS motif undergoes phosphorylation in keratinocytes and is very likely to mediate binding to 14-3-3σ which, as mentioned above, is known to play a role in YAP1 regulation and keratinocyte differentiation (Sambandam et al., 2015; Schlegelmilch et al., 2011). Even though K15 does not feature a SCRAPS motif in its N-terminal head domain, co-IP assays showed that, nevertheless, 14-3-3σ occurs in immunoprecipitates targeted at K15. Unlike 14-3-3σ and K14, however, 14-3-3σ and K15 lack spatial proximity in keratinocytes. From this we infer that K14 and K15 relate differently to 14-3-3σ, certainly at a spatial level, though key details are lacking. Several elements stand out as candidate contributors to the keratin/14-3-3/YAP1 interplay in keratinocytes being programmed to undergo differentiation. First, as we report here, 14-3-3Pred predicts that K5, K14’s preferred co-polymerization partner, is itself a potential 14-3-3 binding protein. One of the high scoring potential sites for 14-3-3 binding on K5, Serine 18 (see Supp. Fig. 4B), is conserved and undergoes phosphorylation in both human and mouse keratinocytes. Accordingly, K5 could be a participant or modulator of the K14/14-3-3σ/YAP1 complex. Second, it is also possible that, at its core, the K14-dependent regulation of YAP1 is not significantly dependent on 14-3-3σ, such that the significance of the SCRAPS motif lies elsewhere. Third, the presence of 14-3-3σ in GFP-K15 immunoprecipitates may reflect interactions with a non-keratin protein present in the complex. Fourth, we previously showed that the cysteine residue located in the SCRAPS motif partakes in K14-dependent disulfide bonding in keratinocytes (Feng and Coulombe, 2015a) – but whether it plays a role in YAP1 regulation has yet to be addressed. Fifth and finally, mechanical cues play a significant role as progenitor keratinocytes commit to differentiation (see Biggs et al., 2020; Miroshnikova et al., 2018; Nekrasova et al., 2018; Ning et al., 2021), and may influence biochemical interactions between keratin, 14-3-3α, YAP1 and other relevant proteins. Future efforts are needed to define when and how, precisely, K14 and other keratins engage 14-3-3 adaptors, the specific role of phosphorylation (and the kinases involved) in these interactions, and how these events relate in space and time to the binding and regulation of YAP1.

How K15 promotes the progenitor state in the epidermis, and whether it may go beyond acting as an antagonist of K14 with regards to the negative regulation of YAP1’s transcriptional role, are unknown. Similarly to K14, the evidence in hand does not support a direct role for K15 in cell cycle control (Cockburn et al., 2022; Cohen et al., 2022). In skin, *KRT15*^high^ keratinocytes occur infrequently in G2/M-phase^high^ or S-phase^high^ subpopulations (this study; also, see Morris et al., 2004; Trempus et al., 2003). Accordingly, we envision that a higher K15:K14 ratio keeps keratinocytes in a progenitor state in which they remain sensitive to proliferation cues. The latter is consistent with the original description of K15 as being enriched in slow-cycling keratinocytes of the hair bulge (Lyle et al., 1998). On the other hand, we report here that the *KRT15* (human) and *Krt15* (mouse) transcripts are, at the single cell level, highly correlated with genes defining the relationship of keratinocytes with the extracellular matrix. Many of these genes are *bona fide* YAP1 targets (e.g., *CYR61*, *CCND2*, *TGFB1*) and/or are known to foster a microenvironment that promotes epithelial stemness (e.g., *COL17A1*, *ITGA1*, *POSTN*) (Morris et al., 2004; Tumbar et al., 2004). We also show that the ultrastructural attributes of the basal lamina, the size of basal keratinocytes and the morphological attributes of their nuclei each are significantly altered in *Krt14^C373A/null^* mice. How such differences arise, molecularly, remains unknown at this point. Previous studies have described a conspicuous lack of keratin bundling and the presence of so-called “wispy” keratin filaments in basal keratinocytes of *KRT14* null human (Chan et al., 1994; Jonkman et al., 1996; Rugg et al., 1994) and *Krt14* null mouse skin (Lloyd et al., 1995). While this may be a function of the lower prevailing levels of K15 protein in such circumstances, our observations of cells transfected with K14 vs. K15 supports the notion that these two keratins promote the formation of filaments having different properties. The different complement of cysteine residues (including the stutter cysteine) in K14 and K15 may play a role in this regard (see Feng and Coulombe, 2015a). The paucity of information about K15 applies not only to its physiological role(s) but also its polymerization and other biochemical properties.

With the notable exception of *KRT15,* initial insight about the functional importance of the keratin genes expressed in the epidermis was inferred from the phenotype of transgenic mouse models and/or individuals suffering from rare genetic skin disorders. The main disorder associated with *KRT14* (and *KRT5*), EBS, is typically caused by dominantly-acting missense alleles (Coulombe et al., 2009) but also arises in individuals whose genome is homozygous for the equivalent of null alleles (Chan et al., 1994; Jonkman et al., 1996; Rugg et al., 1994). No disease association has been reported yet for *KRT15*. Yet, GnomAD (Karczewski et al., 2020) reports variants at several codons in *KRT15*, with many predicted to act deleteriously including alleles altering Arg 115. The latter corresponds to a prominent mutational hotspot in *KRT14* (Arg 125; Coulombe et al., 2009), in other type I keratin genes expressed in epidermis including *KRT10* and *KRT9* (Coulombe and Lee, 2012; Szeverenyi et al., 2008), and is consistently associated with severe clinical presentations. Per GnomAd (Karczewski et al., 2020), Arg 115 in *KRT15* is mutated at an appreciable frequency in the human population and yet no disease association has been reported. Moreover, mice null for *Krt15* are viable and do not exhibit skin fragility (see below) whereas mice null for *Krt14* (Lloyd et al., 1995) or *Krt5* (Peters, 2001) show extensive and penetrant skin and oral blistering, and die shortly after birth. At least three known factors likely account for these striking differences. First, K15 is expressed at lower levels than K14 (and K5) in basal keratinocytes – this is so at the transcript level in human and mouse epidermis (this study) and at the protein level in newborn mouse skin (Feng, 2013) – such that the consequences of its mutated form may be mitigated by the presence of K14 at higher levels. Second, K15 expression is highest in basal cells located in the deeper segments of epidermal rete ridges – these cells may be protected from frictional trauma while, transcriptionally, they exhibit higher expression of genes mediating adhesion to the ECM (this study; see Morris et al., 2004; Tumbar et al., 2004). As a non-exclusive, third possibility, it may be that incorporation of K15 into the keratin network of basal keratinocytes does not enhance mechanical properties as readily as incorporation of K14 does (Lee and Coulombe, 2009) (consistent with this, we found that K15 is more readily detergent-extractable than K14 in transfected HeLa cells; Fig. 6A,B). At present, therefore, *KRT15*/K15 stands out among the main epidermal keratins given a lack of known involvement in a human genetic disease.

Mice in which the *Krt15* locus has been targeted for inactivation have been reported (Cui et al., 2022; Ievlev et al., 2023) but a detailed characterization of their skin phenotype is not available. *Krt15* null mice are viable and, while they do not show any sign of epithelial fragility, they exhibit patchy hair loss as young adults (Cui et al., 2022; Ievlev et al., 2023). The latter could reflect a hair cycle defect and/or a premature exhaustion of the hair bulge-associated stem cell pool, a possibility consistent with our model. While it was focused on studying the trachea, which features a pseudostratified epithelium, a prior effort by levlev *et al*. (Ievlev et al., 2023) yielded findings that directly support our model. levlev *et al*. showed that mice targeted at the *Krt15* locus exhibit impaired basal cell proliferation in the airways but no alteration in their subsequent differentiation. By contrast, mice targeted at the *Krt14* locus in a similar manner show enhanced proliferation of basal cells along with impaired differentiation into ciliated and club cell types. These authors also observed a decrease in label-retaining basal cells in the tracheal epithelium at 21 days after injury in *Krt15* null mice. Such findings are, again, entirely consistent with ours and provide strong support for our model. In conceptually distinct studies, the Tumbar laboratory showed that, relative to that of *Krt14*, expression of *Krt15* is stronger in rete ridges of human skin and in the “scale” domain of healthy mouse tail skin (Ghuwalewala et al., 2022). This correlates with local and stable differences in proliferation rate (higher in K15-enriched rete ridges and mouse tail scales), K10 expression and, ultimately, in differentiation lineages being pursued (Ghuwalewala et al., 2022; Sada et al., 2016). These findings are again consistent with our model stating that a higher K15:K14 ratio promotes the progenitor state but is neutral with regards to other cues regulating proliferation rates and differentiation lineages.

## MATERIALS AND METHODS

### Mouse studies

All mouse studies were reviewed and approved by the Institutional Animal Use and Care Committee (IACUC) at the University of Michigan. All mice (C57BL/6 strain background) were maintained under specific pathogen-free conditions, fed rodent chow and water ad libitum, and kept under a 12-h day/night cycle. Male and female C57Bl/6 mice of 2–3 months of age (young adults) were used for all studies unless indicated otherwise. *Krt14^C373A/C373A^*mice were previously reported in (Guo et al., 2020). Cryopreserved sperm from *Krt14^WT/null^* mice was purchased from MMRRC (Strain name: C57BL/6N-Krt14 /MbpMmucd tm1.1(KOMP)Vlcg; RRID:MMRRC_048362-UCD). In vitro fertilization of eggs harvested from C57Bl/6 females was performed by the University of Michigan Transgenic Core. Pups were screened for the null allele using standard PCR assays with oligonucleotides listed in STAR*Methods, and a transgenic line was established and maintained (C57BL/6 background) through standard husbandry methods.

### Alignment of K14 and K15 amino acid sequences and 14-3-3 binding site prediction

Sequence alignments for human and mouse K14 and K15 were done using the Clustal Omega freeware in the UniProt alignment tool. Computational predictions for 14-3-3 binding were carried out using 14-3-3pred (https://www.compbio.dundee.ac.uk/1433pred) (Madeira, 2015). Lollipop plots were generated using R.

### YAP1 indirect immunofluorescence and quantitation of nuclear:cytoplasmic signal ratio

HeLa cells were purchased from ATCC and were routinely tested for mycoplasma using the MycoAlert® Mycoplasma Detection Kit (Lonza, LT07-118). Cells were cultured in 4.5 g/L D-glucose DMEM (Gibco, 11995073) supplemented with 1% penicillin and streptomycin (Gibco, 15140122) and 10% fetal bovine serum. TWIST mammalian expression plasmids were synthesized by Twist Biosciences (San Francisco, CA). Codon-optimized *KRT14* and *KRT15* cDNAs were synthesized into a pTwist-CMV vector containing a CMV enhancer and promoter for expression in mammalian cells. All plasmids were verified by whole plasmid sequencing (Plasmidsaurus). HeLa cells were transfected using the SE Cell Line 4D-Nucleofector kit (Lonza, V4XC-1032) and program DS-138. 300,000 cells and a total of 2 μg of plasmid DNA were transfected per parameter. After transfection, cells were plated on #1.5 glass coverslips and cultured for 24 h. After 24 h, media was removed, and cells were fixed in 4% PFA/PBS for 10 min. Fixed cells were washed, permeabilized for 5 min in 0.1% Triton X-100, and then incubated for 1 h with rabbit anti-YAP1 antibody (Cell Signaling Technology, 14074S), followed by 1 h incubated with an Alexa Fluor conjugated secondary antibody, and counterstained with DAPI. Coverslips were mounted on a slide with Fluorsave (Millipore, 345789). Coverslips were imaged with a 63X objective with a Zeiss LSM 800 confocal microscope. Laser intensity and detector gain were optimized for each fluor/channel. YAP1 localization was quantified using Zeiss ZEN Lite software. Using the spline tool, the cytoplasm (defined by the mCherry channel) and the nucleus (defined by the DAPI channel) were traced. The area and mean intensity values (MIV) for each channel were calculated for the cytoplasmic and nuclear objects. The cytoplasmic object also contains the nuclear object. The sum intensity value (SIV) of YAP1 was calculated by multiplying the area by the mean intensity value for the cytoplasmic and nuclear objects. The nuclear YAP1 sum intensity value was subtracted from the cytoplasmic YAP1 sum intensity value. The MIV of the resultant nuclear subtracted cytoplasmic SIV was calculated by dividing by the nuclear area subtracted from the cytoplasmic area. The Nuclear:Cytoplasmic YAP1 ratio was calculated for each cell and controlled for transfection by correction and normalization for 488 channel intensity. Statistical analysis was performed in GraphPad Prism. Conditions were compared with Mann-Whitney tests. For figure visualization, datapoints were normalized to the mean of the “K5/no plasmid” sample condition.

### YAP1-14-3-3σ proximity ligation assay (PLA) and quantitation

pMAX-GFP was supplied from Lonza. All keratin overexpression constructs were synthesized by Twist Biosciences and subjected to whole plasmid sequencing (Plasmidsaurus). HeLa cells were transfected using SE Cell Line 4D-Nucleofector kit (Lonza, V4XC-1032) and program CN-114. 150,000 cells and 1 ug of total plasmid DNA were transfected per parameter. After transfection, cells were plated on #1.5 glass coverslips and cultured for 24 h. After 24 h, media was removed, and cells were rinsed with 1X PBS and fixed in 4% PFA/PBS for 10 min. Fixed cells were washed, permeabilized for 10 min in 0.1% Triton X-100, and then blocked in 2.5% NDS/PBS overnight at 4C. Cells were then incubated for 1 h at 37C with rabbit anti-YAP1 antibody (Cell Signaling, 14074S) and goat anti-14-3-3σ antibody (Abcam, ab77187). Following primary antibody incubation, cells were incubated with anti-rabbit PLUS and anti-goat MINUS DuoLink Probes (Sigma-Aldrich, DUO92002, DUO92006), and PLA signal was developed according to manufacturer protocol (Sigma-Aldrich, DUO92013). Coverslips were imaged with a 40X objective with a Zeiss LSM 800 confocal microscope. Laser intensity and detector gain were optimized for each fluor/channel. Images were taken as Z-stacks spanning 10µm at 1µm intervals. ImageJ was used to generate maximum intensity projection images, and PLA punctae/cell were quantified using the ImageJ multi-point tool, using GFP autofluorescence to define the boundaries of the cell(s).

### 14-3-3σ-GFP co-immunoprecipitation

HeLa cells were lysed and diluted according to manufacturer’s direction for GFP-Trap Magnetic Agarose. Briefly, cells were lysed in a solution of 10 mM Tris/Cl pH 7.5, 150 mM NaCl, 0.5 mM EDTA, and 0.5 % Nonidet™ P40 Substitute, and diluted in a 10 mM Tris/Cl pH 7.5, 150 mM NaCl, 0.5 mM EDTA solution. GFP-Trap Magnetic Agarose (Proteintech, #gtma-100) was used, and 25 μl of beads were used, according to the manufacturer’s protocol. Whole cell lysates (600 μg of total protein) were incubated with beads at 4°C for 1hr. The eluted IP samples were denatured and subjected to Western blotting.

### Quantification of 14-3-3σ pulldown efficiency

The Enrichment Ratio described in Fig. 4B’ was calculated as follows:

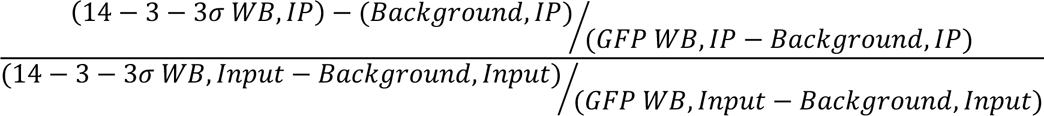

Wherein WB is western blot intensity, as measured via ImageJ.

### RT-qPCR

For tissue, RNA was isolated using TriZOL (Fisher Scientific, #15596018) according to manufacturer’s protocol. For cells, RNA was isolated using Qiagen RNeasy mini kit (Qiagen #74104) following the manufacturer’s protocol. Total RNA from either tissue or cells was converted to complementary DNA (cDNA) using iScript cDNA Synthesis Kit (Bio-Rad #1708891). The cDNA obtained was subjected to qRT-PCR using the iTaq Universal SYBR green kit (Bio-Rad, #1725122) and the CFX 96 Real-Time System (Bio-Rad). The PCR parameters for qRT-PCR were 95°C for 5 min, followed by 40 cycles of 95°C for 10 s and 56°C for 30 s. A “no cDNA template control” and a “melt curve” were included in every PCR run. The normalized expression value of the target gene was determined by first averaging the relative expression of the target gene for each cDNA sample (ΔCq = average Cqtarget gene—average Cqreference gene) and then normalizing the relative expression value of the experimental condition to the control condition (2-(ΔCqExperimental-ΔCqControl)). Primers used in qRT-PCR assays are listed in Supplementary Table X.

### Human buttocks skin indirect immunofluorescence

Sections of paraformaldehyde-fixed, paraffin-embedded healthy human buttocks skin was provided by Dr. Johann Gudjonsson’s laboratory (Department of Dermatology, University of Michigan Medical School) as part of a IRB-approved protocol (see (Cohen et al., 2024)). Sections were deparaffinized in Histo-Clear (National Diagnostics, HS-200), and rehydrated in (2x) 100% EtOH, 95% EtOH, 70% EtOH, and running distilled water. Slides were blocked in 2% normal donkey serum/1% BSA/PBS for 1 hour at RT, incubated in primary antibody diluted in blocking solution (1:200, chicken anti-keratin 14 (BioLegend, 906004); 1:200 chicken anti-keratin 15 (BioLegend, 833904); 1:200 mouse anti-keratin 14 (Abcam, ab80524); 1:200 rabbit anti-keratin 10 (BioLegend, 905404)) for 16 h at 4 °C, Alexa Fluor-conjugated secondary antibody for 1 h at 37 °C, counterstained with DAPI, and mounted in Fluorsave (Millipore, 345789) and a #1.5 cover-slip. Rabbit anti-Ki67 (1:200, Cell Signaling Technology, 9129) indirect immunofluorescence was performed as above with an additional antigen retrieval step following rehydration with running distilled water: slides were gently simmered in 1 mM EDTA, pH 8.0, for 10 minutes, and cooled to room temperature in 1X PBS. Slides were imaged with a 40X objective with a Zeiss LSM 800 confocal microscope. Laser intensity and detector gain were optimized for each fluor/channel. Scale bars represent 50 µm.

### Isolation of newborn skin keratinocytes for primary culture

Newborn mouse keratinocytes were harvested from P0, P1, and P2 pups and seeded in primary culture as follows (see (Guo et al., 2020)). Pups were decapitated, sterilized in 10% (w/v) Povidone-Iodine Solution (Fisher Scientific, 3955-16), and rinsed in 70% EtOH and distilled water. Back skin was dissected and suspended for 18 h in 0.25% Trypsin-EDTA (Gibco, 25200056) at 4 °C. The epidermis was then removed from the dermis and placed basal layer side up in a cell culture dish lid. The living keratinocytes were carefully scraped from the waxy cornified layer and resuspended in basal medium (calcium-free DMEM (US Biological, D9800-10, 70% v/v), Ham’s F-12 (Gibco, 11765054, 22% v/v), Chelex-treated FBS (Bio-Rad, 1421253, 10% v/v), adenine (Sigma-Aldrich, A8626-5G, 0.18 mM), hydrocortisone (Sigma-Aldrich, H4001-1G, 0.5 ug/mL), insulin (Sigma-Aldrich, I9278, 5 ug/mL), cholera toxin (ICN, 856011, 10 nM), EGF (Sigma-Aldrich, E4127, 10 ng/mL), GlutaMAX (Gibco, 35050061, 2 mM), sodium pyruvate (Gibco, 11360070, 1 mM), penicillin (Sigma-Aldrich, P3032-1MU, 100 U/mL)). Pooled back skin keratinocytes were then layered on top of Lymphoprep (Stem Cell Technologies, 07851) and spun for 30 min at 800 x g. The top layer of basal media, containing living cells, was then removed and respun to pellet. The pelleted cells were resuspended in fresh basal media, plated onto #1.5 coverslips, and cultured for up to 72 h. For fixation, media was removed from cells, replaced with 4% PFA/PBS for 10 min, followed by 4 rinses with 1X PBS.

### Transfection and calcium switching of mouse skin keratinocytes in primary culture

Primary cultures of keratinocytes were established as described above. Prior to plating, skin keratinocytes were transfected using P3 Primary Cell 4D-Nucleofector™ X Kit (Lonza, V4XP-3032) and program DS-138. Codon optimized *KRT14* and *KRT15* cDNA was synthesized into a EGFP-C3 vector containing a CMV enhancer and promoter for expression in mammalian cells Twist Biosciences; see above). 0.5 ug of DNA was transfected per parameter. After transfection, cells were plated on #1.5 glass coverslips and cultured for 48 h. Cells were then calcium differentiated by removing media and replacing with basal media supplemented with calcium chloride (Sigma-Aldrich, C7902, 1.2 mM) for 24 h. Cells were fixed by removing the media and replacing with 4% PFA/PBS for 10 min, then rinsed 4x with 1X PBS.

### Indirect immunofluorescence for nuclear YAP1 and its quantitation

Transfected keratinocytes in primary culture were fixed, washed, permeabilized for 5 min in 0.1% Triton X-100, and incubated in primary antibody diluted in 5% NDS/1X PBS blocking buffer (1:100, rabbit anti-YAP1 (Cell Signaling Technology, 14074)) for 1 h at 37 °C, followed by 1 h at 37 °C in an Alexa Fluor conjugated secondary antibody diluted in PBS, and counter-stained with DAPI. Coverslips were mounted on a slide with Fluorsave (Millipore, 345789). Coverslips were imaged using a 40X objective fitted on a Zeiss LSM 800 confocal microscope. Laser intensity and detector gain were optimized for each fluor/channel. YAP1 intensity mean value was quantified using Zeiss ZEN Lite software. Using the spline tool, the nuclei of EGFP-keratin fluorescing cells was traced and the YAP1 mean intensity value was calculated.

### Indirect immunofluorescence for K14 and K15 on primary keratinocytes and quantitation

Fixed cells were washed, permeabilized for 5 min in 0.1% Triton X-100, and then incubated in primary antibody diluted in 5% NDS/1X PBS blocking solution (1:200, chicken anti-keratin 14 (BioLegend, 906004); 1:200 chicken anti-keratin 15 (BioLegend, 833904)) for 1 h at 37 °C, followed by 1 h at 37 °C in an Alexa Fluor conjugated secondary antibody diluted in PBS, and finally counterstained with DAPI. Coverslips were mounted on a slide with Fluorsave (Millipore, 345789). Coverslips were imaged with a 63X objective with a Zeiss LSM 800 confocal microscope. Laser intensity and detector gain were optimized for each fluor/channel. K14 and K15 intensity mean value were quantified using Zeiss ZEN Lite software. Using the spline tool, the cytoplasm was traced and the mean intensity values for each channel were calculated. There were no apparent alterations to keratin filament morphology, emulating previous observations in *Krt14^C373A/C373A^* primary keratinocytes (Guo et al., 2020).

### Seurat clustering

Single cell RNAseq datasets from human trunk skin and foreskin were originally published by Cheng et al. (Cheng et al., 2018). The data was reanalyzed using Seurat’s SCTransform “v2” as detailed in a previous publication (Cohen et al., 2024). Analysis For the single cell RNAseq dataset from saline-treated mouse back skin (GSE194254) (Leyva-Castillo et al., 2022) were done as follows: cells were filtered to remove doublets using scDblFinder (Germain et al., 2021), percent mitochondrial genes (<15%) and number of features (400< x <4000). Retained cells (12,093 cells) were then processed using harmony to integrate 20 principal components and perform dimensional reduction, with 0.5 resolution for clustering. This analysis resulted in the identification of 22 unique clusters. Enrichr (Kuleshov et al., 2016) was used to analyze the Seurat clusters for keratinocyte gene signatures using PanglaoDB dataset (Franzen et al., 2019), identifying 7 keratinocyte cell clusters used for further analysis (4,806 cells). Data clustering and analysis was performed using R version 4.4.1, and Seurat version 5.

### Generation of Transcription Signature Composite Scores

Composite scores (Differentiation score, Basal score, G2M Score, S Score) were calculated using arithmetic mean of Seurat normalized natural log values. The following genes (and their direct mouse orthologues) were included in composite scores as follows:

Basal score: *POSTN, ITGB1, LAMA5, COL17A1,DST.*
Differentiation score: *DSP, DMKN, KRTDAP, DSG1, SBSN. G2M*
Score: *MKI67, TOP2A, CDC20, BUB3, CCNB2*.
S Score: *RPA2, UHRF1, ATAD2, RFC2, RRM2*

These scores have been previously shown to accurately represent the cellular states and transitions across epidermal keratinocyte differentiation (Cohen et al., 2022; Cohen et al., 2024). A difference score between basal and differentiation (‘Differentiation Score -Basal Score’) was used to organize keratinocyte clusters according to their natural continuum when analyzing keratin level per cluster (Figure2, Supp. Figure2 and Supp. Figure3). Last, expression of individual keratin genes *(KRT14*, *KRT15*, *KRT10*, *KRT2*), was analyzed against composite scores on a Seurat cluster or single-cell level. Graphs and statistical analysis were generated using Graphpad Prism 9.1.0, and the statistical methods used are reported within figure panels.

### Pup observations, weights, and histological analysis

Pups born from crosses of *Krt14^C373A/C373A^* and *Krt14^WT/null^*were observed daily for 3 weeks. In the first postnatal week, pups were not touched. Observations were performed through the cage so as not to disturb the breeding pair. After week 1, the nest was moved to take pictures of the pups. Between weeks 2 and 3, pups were weighed daily in the afternoons in a weigh boat using a small scale. Pups were tracked based on unique identifying tail markings. For histological analysis, pups were euthanized via decapitation. Back skin was harvested and submerged overnight in an excess of 10% neutral buffered formalin. After fixation, tissue was arranged into cassettes, stored in 70% EtOH at RT, and handed to the University of Michigan Department of Pathology Research Histology Core for paraffin embedding and sectioning. FFPE tissue was deparaffinized in Histo-Clear (National Diagnostics, HS-200), and rehydrated in (2x) 100% EtOH, 95% EtOH, 70% EtOH, and running water. Hematoxylin (Thermo Scientific, TA-060-MH) and eosin Y (Thermo Scientific, 71204) staining was performed per Sigma-Aldrich Procedure No. MHS #1. Epidermal thickness was quantified in ImageJ by measuring from bottom of the basal layer to the top of the granular layer perpendicular to the basal lamina at the point. Measurements were collected every 15 µm of basal lamina. Hair follicle length was quantified in ImageJ by measuring from the bottom of the hair follicle bulb to the basal layer of the infundibulum. Only hair follicles with visible bulbs were measured. Statistical analysis was performed in GraphPad Prism. Conditions were compared with Mann-Whitney tests.

### Dot blot analysis of skin tissue proteins

Tail skin (roughly 1 inch in length, towards the base of the tail) was harvested from 8 week old, male and female, *Krt14^C373A/WT^*and *Krt14^C373A/null^* littermates. Skin was finely minced with fresh razorblades and nutated for 24 h at 4 °C in urea lysis buffer (pH 7.0, 6.5M urea (Sigma-Aldrich, U5378), 50 mM Tris-HCl, 150 mM sodium chloride, 5 mM ethylenediaminetetraacetic acid (EDTA), 0.1% Triton X-100, cOmplete Proteinase Inhibitor Cocktail (Roche, 11697498001)). Tissue was then homogenized on ice with a probe homogenizer and spun at 16,200 x g for 30 min at 4 °C. The insoluble pellet was discarded, and protein concentration of the supernatant was determined by BCA assay (Thermo Scientific, 23225). Dot blotting was performed using a 96-well Minifold I Dot-Blot System (Cytiva, 10447900). Samples were eluted through a 0.45 µm nitrocellulose membrane (BioRad, 1620115) overlaying thick pure cellulose chromatography paper (Fisher Scientific, 05-714-4). Membrane and paper were pre-wet with 3 washes of PBS. Wells were preloaded with 200 µL of 0.01% BSA/PBS. 7.5, 5, 2.5, 1, and 0.5 µg of lysate was added directly to 0.01% BSA/PBS. Samples were eluted and vacuum remained on for 5 min to dry the membrane. Membranes were blocked in 5% BSA/PBS for 1 h following transfer. After blocking, membranes were cut and incubated in primary antibody (1:1000 rabbit anti-histone H3 (Abcam, ab1791); 1:1000 chicken anti-keratin 14 (BioLegend, 906004); 1:1000 chicken anti-keratin 15 (BioLegend, 833904)) diluted in 5% BSA/PBS blocking buffer overnight at 4 °C. Membranes were washed in PBS-T, incubated for 1 h at RT in HRP-conjugated secondary antibodies, and washed prior to visualization with ECL (Cytiva, RPN2232). Mean gray values (MGV) of dots were quantified using ImageJ elliptical selection tool and measurement function. K14 and K15 MGV were normalized for loading by dividing by histone H3 MGV. For data visualization, MGV of each datapoint (corresponding to a single dot) were divided by the mean of the *Krt14^C373A/WT^* populations.

### Histological analysis and quantification of basal keratinocyte density

At 8-weeks of age, male and female *Krt14^C373A/null^* animals appeared healthy and were indistinguishable from *Krt14^C373A/WT^* control littermates. Tissue was harvested from the tail, ear, and tongue and immediately frozen at -40 °C in Tissue-Tek OCT compound (Sakura, 4583). Hematoxylin (Thermo Scientific, TA-060-MH) and eosin Y (Thermo Scientific, 71204) staining was performed on fresh, frozen tissue. 5 µm sections were placed onto slides and air dried for 10 min. Hematoxylin and eosin Y staining was performed per Sigma-Aldrich Procedure No. MHS #1. Basal cell crowding was quantified on skin sections of paraformaldehyde-fixed, paraffin embedded (FFPE) tissue. Tail skin was dissected from 8 week old, male and female, *Krt14^C373A/WT^* and *Krt14^C373A/null^* littermates. Harvested tissue was fixed for 18 h at 4 °C in 10% neutral buffered formalin (Sigma-Aldrich, HT501128). After fixation, tissue was arranged into cassettes, stored in 70% EtOH at RT, and handed to the University of Michigan Department of Pathology Research Histology Core for paraffin embedding and sectioning. FFPE tissue was deparaffinized in Histo-Clear (National Diagnostics, HS-200), and rehydrated in (2x) 100% EtOH, 95% EtOH, 70% EtOH, and running water. Slides were blocked in 5% normal donkey serum/1% BSA/PBS for 1 h at RT, incubated in primary antibody (1:200, chicken anti-keratin 14 (BioLegend, 906004); 1:200, chicken anti-keratin 15 (BioLegend, 833904)) for 16 h at 4 °C, Alexa Fluor-conjugated secondary antibody for 1 h at RT, counterstained with DAPI, and mounted in Fluorsave (Millipore, 345789) and a #1.5 coverslip. Slides were imaged with a 40X objective with a Zeiss LSM 800 confocal microscope. Laser intensity and detector gain were optimized for each fluor/channel. Area was calculated by tracing individual basal keratinocytes. The cytoplasm was distinguished by K15 immunofluorescence staining. K15 is a suitable for this purpose as it is a pan-cytoplasmic stain with distinct gaps at cell peripheries. To calculate basal cell density, basal lamina distance was measured and cells along this distance were counted. Statistical analysis was performed in GraphPad Prism. Conditions were compared with Mann-Whitney tests.

### Mouse tail skin YAP1 and Ki67 indirect immunofluorescence and quantitation

Tail skin was dissected from 8-week-old male and female *Krt14^C373A/WT^* and *Krt14^C373A/null^* tail skin processed for FFPE, and immunostained, as described above. Slides were blocked in 5% normal donkey serum/1% BSA/PBS for 1 h at RT, incubated in primary antibody (1:200 rabbit anti-Ki67 (Cell Signaling Technology, 9129S); 1:200 chicken anti-keratin 15 (BioLegend, 833904); 1:100 rabbit anti-YAP1 (Cell Signaling Technology, 14074)) for 16 h at 4 °C, Alexa Fluor-conjugated secondary antibody for 1 h at room temperature, counterstained with DAPI, and mounted in Fluorsave (Millipore, 345789) and a #1.5 coverslip. Slides were imaged with a 40X objective with a Zeiss LSM 800 confocal microscope. To quantify Ki67 intensity and positivity, the nucleus of individual basal keratinocytes was traced in ZEISS ZEN Lite using the spline tool. The nucleus was distinguished by DAPI immunofluorescence staining. The intensity mean values for each channel per traced cell were then measured. Statistical analysis (Mann-Whitney test) and visualization were performed using GraphPad Prism. The mean Ki67 intensity (arbitrary units) was compared for the entire population of nuclei between *Krt14^C373A/WT^*and *Krt14^C373A/null^* samples. To calculate the percentage of Ki67 positive nuclei, a threshold for Ki67 positivity was set by selecting a population of nuclei with the visually lowest, to absence of, Ki67 signal. The median Ki67 intensity of this population was determined to be 7000 arbitrary units. The percentage of cells per genotype exceeding this “Ki67 positive threshold” was calculated.

### Transmission electron microscopy and quantitation

Tail, ear, and back skin were harvested from 8-week-old, *Krt14^C373A/WT^*and *Krt14^C373A/null^*, male and female littermates. Back skin was shaved prior to dissection. Harvested tissue was pinned to a wax block and initially fixed for 5 min in ice cold 3% glutaraldehyde/3% paraformaldehyde/0.1 M sodium cacodylate (Electron Microscopy Sciences, 15950). Flat, partially fixed tissue was further trimmed into small pieces using fresh razorblades. Subsections were dissected along the sagittal plane of the animal. Tissue pieces were submerged in ice cold 3% glutaraldehyde/3% paraformaldehyde/0.1 M sodium cacodylate and stored at 4 °C until further processing, which was performed by the Michigan Medicine Microscopy Imaging Laboratory for embedding and sectioning. Tissue pieces were sequentially post-fixed in 1.5% potassium ferrocyanide and 2% osmium tetroxide, extensively washed in buffer and distilled water in-between steps, en-bloc stained in 2% uranyle acetate, dehydrated in graded series of ethanol (up to 100%) and pure acetone, and embedded in Durcupan epoxy resin using standard procedures. 70-nm thick sections were cut on an ultramicrotome equipped with a diamond knife and imaged using a JEOL JEM 1400 Plus transmission electron microscope. To calculate basal lamina convolution, we traced the basal lamina to measure the distance processed and divided this by the distance covered by a vector connecting the beginning and end of the traced basal lamina.

### Analysis of Keratin Phosphorylation by Mass Spectrometry

Pelleted N-TERT keratinocytes were lysed in 6.5 M Urea, 50 mM Tris-HCl pH 7.0, 150 mM sodium chloride, 5 mM ethylenediaminetetraacetic acid (EDTA), 0.1% Triton X-100, 50 μM N-ethylmaleimide supplemented with Protease Inhibitor (A32963, Thermo Fisher Scientific) and Phospho-stop phosphatase inhibitor tablets (A32957, Thermo Fisher Scientific). After ultrasonic homogenizing, the lysate was centrifuged at 15,000xg for 10 minutes at 4°C and the cleared supernatants were loaded onto an SDS-PAGE gel. A gel piece of around 50kDa was cut from the Coomassie-stained gel and proceeded with enzymatic digestion using Trypsin/Lys-C mix (PAV5073, Promega) after reduction with 10 mM Dithiothreitol and alkylation with 50 mM 2-Chloroacetamine (C0267, Sigma-Aldrich). The resulting peptides were extracted and cleaned using C18 and dried in speed-vac. The samples were processed and analyzed at the Mass Spectrometry Facility of the Department of Pathology at the University of Michigan. Around 1 mg of peptides was reconstituted in 5% Formic Acid and analyzed in an Orbitrap Ascend Tribrid mass spectrometer (Thermo Fisher Scientific) equipped with a FAIMS unit. The samples were analyzed in a 60 min gradient using two CVs (-45 and -65V). The full scans (MS1) were acquired over a mass-to-charge (m/z) range of 375-1450 with positive polarity and were allowed for detection in data-dependent acquisition (DDA) mode. For MS1 spectra acquisition, the resolution was set to 120,000, the automatic gain control (AGC) target was 1e6, and the maximum injection time was 251 ms. MS2 spectra acquisition was performed at a resolution of 15,000, with an AGC target of 5e4, a maximum injection time of 27 ms. The isolation window was set to 1.6 m/z, collision energy was set to 25%, dynamic exclusion was 60 seconds, and charge exclusion was unassigned, 1, >6. MS data were processed using Fragpipe v.22.0 (Kong et al., 2017) against the Homo sapiens UniProt FASTA database (UP000005640). LFQ-phospho workflow was selected. Enzyme specificity was set to trypsin with up to one missed cleavage. The search included cysteine carbamidomethylation as a fixed modification and oxidation of methionine; N-terminal acetylation; serine, threonine and tyrosine phosphorylation as variable modifications. 1% FDR and >0.6 Phosphorylation Localization score was selected.

## Supporting information

Redmond et al. Suppl. FIGs 1-5 & legends

## ACKNOWLEDGMENTS

The authors are grateful to members of the Coulombe laboratory for advice, to Beau Su for logistical support, and to Venkatesha Basrur (Proteomics Resource Facility), Anna LaForest and Corey Ziebell (Transgenic Core Facility), and Woo Jung Cho and Devon Leroux (Microscopy Imaging Laboratory) at the Univ. Michigan Medical School for technical assistance. CJR (F99 CA274517), SNS (F31 AR038249) and EC (T32 AR007197) received fellowship awards from the National Institutes of Health. EC also received fellowship support from the National Psoriasis Foundation. These studies were supported by grant R01 AR079418 to PAC from the NIH.

## REFERENCES

Biggs, L.C., C.S. Kim, Y.A. Miroshnikova, and S.A. Wickstrom. 2020. Mechanical Forces in the Skin: Roles in Tissue Architecture, Stability, and Function. J. Invest. Dermatol. 140:284–290.

Blanpain, C., and E. Fuchs. 2009. Epidermal homeostasis: a balancing act of stem cells in the skin. Nat. Rev. Mol. Cell Biol. 10:207–217.

Bonifas, J.M., A.L. Rothman, and E.H. Epstein, Jr. 1991. Epidermolysis bullosa simplex: evidence in two families for keratin gene abnormalities. Science. 254:1202–1205.

Bose, A., M.T. Teh, I.C. Mackenzie, and A. Waseem. 2013. Keratin 15 as a biomarker of epidermal stem cells. Int J Mol Sci. 14:19385–19398.

Bunick, C.G., and L.M. Milstone. 2017. The X-Ray Crystal Structure of the Keratin 1-Keratin 10 Helix 2B Heterodimer Reveals Molecular Surface Properties and Biochemical Insights into Human Skin Disease. J Invest Dermatol. 137:142–150.

Chan, Y., I. Anton-Lamprecht, Q.C. Yu, A. Jackel, B. Zabel, J.P. Ernst, and E. Fuchs. 1994. A human keratin 14 "knockout": the absence of K14 leads to severe epidermolysis bullosa simplex and a function for an intermediate filament protein. Genes Dev. 8:2574–2587

Cheng, J.B., A.J. Sedgewick, A.I. Finnegan, P. Harirchian, J. Lee, S. Kwon, M.S. Fassett, J. Golovato, M. Gray, R. Ghadially, W. Liao, B.E. Perez White, T.M. Mauro, T. Mully, E.A. Kim, H. Sbitany, I.M. Neuhaus, R.C. Grekin, S.S. Yu, J.W. Gray, E. Purdom, R. Paus, C.J. Vaske, S.C. Benz, J.S. Song, and R.J. Cho. 2018. Transcriptional Programming of Normal and Inflamed Human Epidermis at Single-Cell Resolution. Cell Reports. 25:871–883.

Cockburn, K., K. Annusver, D.G. Gonzalez, S. Ganesan, D.P. May, K.R. Mesa, K. Kawaguchi, M. Kasper, and V. Greco. 2022. Gradual differentiation uncoupled from cell cycle exit generates heterogeneity in the epidermal stem cell layer. Nat Cell Biol. 24:1692–1700.

Cohen, E., C. Johnson, C.J. Redmond, R.R. Nair, and P.A. Coulombe. 2022. Revisiting the significance of keratin expression in complex epithelia. J Cell Sci. 135.

Cohen, E., C.N. Johnson, R. Wasikowski, A.C. Billi, L.C. Tsoi, J.M. Kahlenberg, J.E. Gudjonsson, and P.A. Coulombe. 2024. Significance of stress keratin expression in normal and diseased epithelia. iScience. 27:108805.

Coulombe, P.A., Hutton, M. E., Letai, A., Hebert, A., Paller, A. S., Fuchs, E. 1991. Point mutations in human keratin 14 genes of epidermolysis bullosa simplex patients: genetic and functional analyses. Cell. 66:1301–1311.

Coulombe, P.A., M.L. Kerns, and E. Fuchs. 2009. Epidermolysis bullosa simplex: a paradigm for disorders of tissue fragility. J Clin Invest. 119:1784–1793.

Coulombe, P.A., R. Kopan, and E. Fuchs. 1989. Expression of keratin K14 in the epidermis and hair follicle: insights into complex programs of differentiation. J. Cell Biol. 109:2295–2312

Coulombe, P.A., and C.H. Lee. 2012. Defining keratin protein function in skin epithelia: epidermolysis bullosa simplex and its aftermath. J Invest. Dermatol. 132:763–775.

Cui, J., Q. Zhao, Z. Song, Z. Chen, X. Zeng, C. Wang, Z. Lin, F. Wang, and Y. Yang. 2022. KLHL24-Mediated Hair Follicle Stem Cells Structural Disruption Causes Alopecia. J Invest Dermatol. 142:2079–2087 e2078.

Feng, X., and P.A. Coulombe. 2015a. Complementary roles of specific cysteines in keratin 14 toward the assembly, organization, and dynamics of intermediate filaments in skin keratinocytes. J Biol Chem. 290:22507–22519.

Feng, X., and P.A. Coulombe. 2015b. A role for disulfide bonding in keratin intermediate filament organization and dynamics in skin keratinocytes. J. Cell Biol. 209:59–72.

Feng, X., Zhang, H., Margolick, J. B., Coulombe, P. A. 2013. Keratin intracellular concentration revisited: implications for keratin function in surface epithelia. J. Invest. Dermatol. 133:850–853.

Franzen, O., L.M. Gan, and J.L.M. Bjorkegren. 2019. PanglaoDB: a web server for exploration of mouse and human single-cell RNA sequencing data. Database (Oxford). 2019.

Fuchs, E. 1995. Keratins and the skin. Ann. Rev. Cell Dev. Biol. 11:123–153.

Fuchs, E., and H. Green. 1980. Changes in keratin gene expression during terminal differentiation of the keratinocyte. Cell. 19:1033–1042.

Germain, P.L., A. Lun, C. Garcia Meixide, W. Macnair, and M.D. Robinson. 2021. Doublet identification in single-cell sequencing data using scDblFinder. F1000Res. 10:979.

Ghuwalewala, S., S.A. Lee, K. Jiang, J. Baidya, G. Chovatiya, P. Kaur, D. Shalloway, and T. Tumbar. 2022. Binary organization of epidermal basal domains highlights robustness to environmental exposure. The EMBO journal. 41:e110488.

Giroux, V., A.A. Lento, M. Islam, J.R. Pitarresi, A. Kharbanda, K.E. Hamilton, K.A. Whelan, A. Long, B. Rhoades, Q. Tang, H. Nakagawa, C.J. Lengner, A.J. Bass, E.P. Wileyto, A.J. Klein-Szanto, T.C. Wang, and A.K. Rustgi. 2017. Long-lived keratin 15+ esophageal progenitor cells contribute to homeostasis and regeneration. J Clin Invest. 127:2378–2391.

Guo, Y., C.J. Redmond, K.A. Leacock, M.V. Brovkina, S. Ji, V. Jaskula-Ranga, and P.A. Coulombe. 2020. Keratin 14-dependent disulfides regulate epidermal homeostasis and barrier function via 14-3-3σ and YAP1. eLife. 9:e53165.

Ievlev, V., T.J. Lynch, K.W. Freischlag, C.B. Gries, A. Shah, A.C. Pai, B.A. Ahlers, S. Park, J.F. Engelhardt, and K.R. Parekh. 2023. Krt14 and Krt15 differentially regulate regenerative properties and differentiation potential of airway basal cells. JCI Insight. 8.

Inoue, K., N. Aoi, T. Sato, Y. Yamauchi, H. Suga, H. Eto, H. Kato, J. Araki, and K. Yoshimura. 2009. Differential expression of stem-cell-associated markers in human hair follicle epithelial cells. Lab Invest. 89:844–856.

Jaks, V., N. Barker, M. Kasper, J.H. van Es, H.J. Snippert, H. Clevers, and R. Toftgard. 2008. Lgr5 marks cycling, yet long-lived, hair follicle stem cells. Nat Genet. 40:1291–1299.

Jonkman, M.F., K. Heeres, H.H. Pas, M.J. van Luyn, J.D. Elema, L.D. Corden, F.J. Smith, W.H. McLean, F.C. Ramaekers, M. Burton, and H. Scheffer. 1996. Effects of keratin 14 ablation on the clinical and cellular phenotype in a kindred with recessive epidermolysis bullosa simplex. J Invest Dermatol. 107:764–769.

Karczewski, K.J., L.C. Francioli, G. Tiao, B.B. Cummings, J. Alfoldi, Q. Wang, R.L. Collins, K.M. Laricchia, A. Ganna, D.P. Birnbaum, L.D. Gauthier, H. Brand, M. Solomonson, N.A. Watts, D. Rhodes, M. Singer-Berk, E.M. England, E.G. Seaby, J.A. Kosmicki, R.K. Walters, K. Tashman, Y. Farjoun, E. Banks, T. Poterba, A. Wang, C. Seed, N. Whiffin, J.X. Chong, K.E. Samocha, E. Pierce-Hoffman, Z. Zappala, A.H. O’Donnell-Luria, E.V. Minikel, B. Weisburd, M. Lek, J.S. Ware, C. Vittal, I.M. Armean, L. Bergelson, K. Cibulskis, K.M. Connolly, M. Covarrubias, S. Donnelly, S. Ferriera, S. Gabriel, J. Gentry, N. Gupta, T. Jeandet, D. Kaplan, C. Llanwarne, R. Munshi, S. Novod, N. Petrillo, D. Roazen, V. Ruano-Rubio, A. Saltzman, M. Schleicher, J. Soto, K. Tibbetts, C. Tolonen, G. Wade, M.E. Talkowski, C. Genome Aggregation Database, B.M. Neale, M.J. Daly, and D.G. MacArthur. 2020. The mutational constraint spectrum quantified from variation in 141,456 humans. Nature. 581:434–443.

Kim, S., P. Wong, and P.A. Coulombe. 2006. A keratin cytoskeletal protein regulates protein synthesis and epithelial cell growth. Nature. 441:362–365.

Kong, A.T., F.V. Leprevost, D.M. Avtonomov, D. Mellacheruvu, and A.I. Nesvizhskii. 2017. MSFragger: ultrafast and comprehensive peptide identification in mass spectrometry-based proteomics. Nature Methods. 14:513–520.

Ku, N.O., Liao, J., Omary, M. B. 1998. Phosphorylation of human keratin 18 serine 33 regulates binding to 14-3-3 proteins. EMBO J. 17:1892–1906.

Kuleshov, M.V., M.R. Jones, A.D. Rouillard, N.F. Fernandez, Q. Duan, Z. Wang, S. Koplev, S.L. Jenkins, K.M. Jagodnik, A. Lachmann, M.G. McDermott, C.D. Monteiro, G.W. Gundersen, and A. Ma’ayan. 2016. Enrichr: a comprehensive gene set enrichment analysis web server 2016 update. Nucleic Acids Research. 44:W90–97.

Lee, C.H., and P.A. Coulombe. 2009. Self-organization of keratin intermediate filaments into cross-linked networks. J. Cell Biol. 186:409–421.

Lee, C.H., M.S. Kim, B.M. Chung, D.J. Leahy, and P.A. Coulombe. 2012. Structural basis for heteromeric assembly and perinuclear organization of keratin filaments. Nat. Struct. Mol. Biol. 19:707–715.

Leyva-Castillo, J.M., L. Sun, S.Y. Wu, S. Rockowitz, P. Sliz, and R.S. Geha. 2022. Single-cell transcriptome profile of mouse skin undergoing antigen-driven allergic inflammation recapitulates findings in atopic dermatitis skin lesions. J Allergy Clin Immunol. 150:373–384.

Li, P., K. Rietscher, H. Jopp, T.M. Magin, and M.B. Omary. 2023. Posttranslational modifications of keratins and their associated proteins as therapeutic targets in keratin diseases. Curr. Opin. Cell Biol. 85:102264.

Lin, Z., S. Jin, J. Chen, Z. Li, Z. Lin, L. Tang, Q. Nie, and B. Andersen. 2020. Murine interfollicular epidermal differentiation is gradualistic with GRHL3 controlling progression from stem to transition cell states. Nat Commun. 11:5434.

Liu, Y., S. Lyle, Z. Yang, and G. Cotsarelis. 2003. Keratin 15 promoter targets putative epithelial stem cells in the hair follicle bulge. J Invest Dermatol. 121:963–968.

Lloyd, C., Q.C. Yu, J. Cheng, K. Turksen, L. Degenstein, E. Hutton, and E. Fuchs. 1995. The basal keratin network of stratified squamous epithelia: defining K15 function in the absence of K14. J Cell Biol. 129:1329–1344.

Lyle, S., M. Christofidou-Solomidou, Y. Liu, D.E. Elder, S. Albelda, and G. Cotsarelis. 1998. The C8/144B monoclonal antibody recognizes cytokeratin 15 and defines the location of human hair follicle stem cells. J Cell Sci. 111 (Pt 21):3179–3188.

Ma, X., H. Zhang, X. Xue, and Y.M. Shah. 2017. Hypoxia-inducible factor 2alpha (HIF-2alpha) promotes colon cancer growth by potentiating Yes-associated protein 1 (YAP1) activity. J Biol Chem. 292:17046–17056.

Madeira, F., Tinti, M., Murugesan, G., Berrett, E., Stafford, M., Toth, R., Cole, C., MacKintosh, C., Barton, G.J.. 2015. 14-3-3-Pred: Improved methods to predict 14-3-3-binding phosphopeptides. Bioinformatics. 31:2276–2283.

Mariani, R.A., S. Paranjpe, R. Dobrowolski, and G.F. Weber. 2020. 14-3-3 targets keratin intermediate filaments to mechanically sensitive cell-cell contacts. Molecular biology of the cell. 31:930–943.

Michel, M., Torok, N., Godbout, M. J., Lussier, M., Gaudreau, P., Royal, A., Germain, L. 1996. Keratin 19 as a biochemical marker of skin stem cells in vivo and in vitro: keratin 19 expressing cells are differentially localized in function of anatomic sites, and their number varies with donor age and culture stage. J Cell Sci. 109:1017–1028.

Miroshnikova, Y.A., H.Q. Le, D. Schneider, T. Thalheim, M. Rubsam, N. Bremicker, J. Polleux, N. Kamprad, M. Tarantola, I. Wang, M. Balland, C.M. Niessen, J. Galle, and S.A. Wickstrom. 2018. Adhesion forces and cortical tension couple cell proliferation and differentiation to drive epidermal stratification. Nat Cell Biol. 20:69–80.

Morris, R.J., Y. Liu, L. Marles, Z. Yang, C. Trempus, S. Li, J.S. Lin, J.A. Sawicki, and G. Cotsarelis. 2004. Capturing and profiling adult hair follicle stem cells. Nat Biotechnol. 22:411–417.

Nekrasova, O., R.M. Harmon, J.A. Broussard, J.L. Koetsier, L.M. Godsel, G.N. Fitz, M.L. Gardel, and K.J. Green. 2018. Desmosomal cadherin association with Tctex-1 and cortactin-Arp2/3 drives perijunctional actin polymerization to promote keratinocyte delamination. Nat Commun. 9:1053.

Ning, W., A. Muroyama, H. Li, and T. Lechler. 2021. Differentiated Daughter Cells Regulate Stem Cell Proliferation and Fate through Intra-tissue Tension. Cell stem cell. 28:436–452 e435.

Pennington, K.L., T.Y. Chan, M.P. Torres, and J.L. Andersen. 2018. The dynamic and stress-adaptive signaling hub of 14-3-3: emerging mechanisms of regulation and context-dependent protein-protein interactions. Oncogene. 37:5587–5604.

Peters, B., Kirfel, J., Bussow, H., Vidal, M., Magin, T. M. 2001. Complete cytolysis and neonatal lethality in keratin 5 knockout mice reveal its fundamental role in skin integrity and in epidermolysis bullosa simplex. Molecular biology of the cell. 12:1775–1789.

Pocaterra, A., P. Romani, and S. Dupont. 2020. YAP/TAZ functions and their regulation at a glance. J Cell Sci. 133.

Porter, R.M., D.P. Lunny, P.H. Ogden, S.M. Morley, W.H. McLean, A. Evans, D.L. Harrison, E.L. Rugg, and E.B. Lane. 2000. K15 expression implies lateral differentiation within stratified epithelial basal cells. Laboratory investigation. 80:1701–1710.

Ramms, L., G. Fabris, R. Windoffer, N. Schwarz, R. Springer, C. Zhou, J. Lazar, S. Stiefel, N. Hersch, U. Schnakenberg, T.M. Magin, R.E. Leube, R. Merkel, and B. Hoffmann. 2013. Keratins as the main component for the mechanical integrity of keratinocytes. Proc Natl Acad Sci U S A. 110:18513–18518.

Rugg, E.L., W.H. McLean, E.B. Lane, R. Pitera, J.R. McMillan, P.J. Dopping-Hepenstal, H.A. Navsaria, I.M. Leigh, and R.A. Eady. 1994. A functional "knockout" of human keratin 14. Genes Dev. 8:2563–2573.

Sada, A., F. Jacob, E. Leung, S. Wang, B.S. White, D. Shalloway, and T. Tumbar. 2016. Defining the cellular lineage hierarchy in the interfollicular epidermis of adult skin. Nat Cell Biol. 18:619–631.

Sambandam, S.A.T., R.B. Kasetti, L. Xue, D.C. Dean, Q. Lu, and Q. Li. 2015. 14-3-3sigma regulates keratinocyte proliferation and differentiation by modulating Yap1 cellular localization. J Invest Dermatol. 135:1621–1628.

Schlegelmilch, K., M. Mohseni, O. Kirak, J. Pruszak, J.R. Rodriguez, D. Zhou, B.T. Kreger, V. Vasioukhin, J. Avruch, T.R. Brummelkamp, and F.D. Camargo. 2011. Yap1 acts downstream of alpha-catenin to control epidermal proliferation. Cell. 144:782–795.

Schweizer, J., Bowden, P. E., Coulombe, P. A., Langbein, L., Lane, E. B., Magin, T. M., Maltais, L., Omary, M. B., Parry, D. A., Rogers, M. A., Wright, M. W. 2006. New consensus nomenclature for mammalian keratins. J. Cell Biol.174:169–174.

Steiner, S.N., E. Horst, M. Athaiya, C.N. Johnson, J.Y. Shen, M.L. Kerns, G. Mehta, R. Iglesias-Bartolome, and P.A. Coulombe. 2025. Under Pressure: A unique mechanoresponsive mechanism of body site-specific keratin regulation in palmoplantar epidermis. bioRxiv.

Stoler, A., Kopan, R., Duvic, M., Fuchs, E. 1988. Use of monospecific antisera and cRNA probes to localize the major changes in keratin expression during normal and abnormal epidermal differentiation. J. Cell Biol. 107:427–446.

Sun, B.K., L.D. Boxer, J.D. Ransohoff, Z. Siprashvili, K. Qu, V. Lopez-Pajares, S.T. Hollmig, and P.A. Khavari. 2015. CALML5 is a ZNF750- and TINCR-induced protein that binds stratifin to regulate epidermal differentiation. Genes Dev. 29:2225–2230.

Szeverenyi, I., A.J. Cassidy, C.W. Chung, B.T. Lee, J.E. Common, S.C. Ogg, H. Chen, S.Y. Sim, W.L. Goh, K.W. Ng, J.A. Simpson, L.L. Chee, G.H. Eng, B. Li, D.P. Lunny, D. Chuon, A. Venkatesh, K.H. Khoo, W.H. McLean, Y.P. Lim, and E.B. Lane. 2008. The Human Intermediate Filament Database: comprehensive information on a gene family involved in many human diseases. Hum Mutat. 29:351–360.

Totaro, A., T. Panciera, and S. Piccolo. 2018. YAP/TAZ upstream signals and downstream responses. Nat Cell Biol. 20:888–899.

Trempus, C.S., R.J. Morris, C.D. Bortner, G. Cotsarelis, R.S. Faircloth, J.M. Reece, and R.W. Tennant. 2003. Enrichment for living murine keratinocytes from the hair follicle bulge with the cell surface marker CD34. J Invest Dermatol. 120:501–511.

Tumbar, T., G. Guasch, V. Greco, C. Blanpain, W.E. Lowry, M. Rendl, and E. Fuchs. 2004. Defining the epithelial stem cell niche in skin. Science. 303:359–363.

Tzivion, G., Z.J. Luo, and J. Avruch. 2000. Calyculin A-induced vimentin phosphorylation sequesters 14-3-3 and displaces other 14-3-3 partners in vivo. J Biol Chem. 275:29772–29778.

Wang, J., Y. Qiu, Y. Zhu, X. Ren, X. Zhou, X. Wang, H. Song, J. Li, C. Gao, G. Zhou, and P. Cao. 2025. Generation of the Krt24-Cre(ERT2) Mouse Line Targeting Outer Bulge Hair Follicle Cells. Int J Mol Sci. 26.

Waseem, A., B. Dogan, N. Tidman, Y. Alam, P. Purkis, S. Jackson, A. Lalli, M. Machesney, and I.M. Leigh. 1999. Keratin 15 expression in stratified epithelia: downregulation in activated keratinocytes. J Invest Dermatol. 112:362–369.

Webb, A., A. Li, and P. Kaur. 2004. Location and phenotype of human adult keratinocyte stem cells of the skin. Differentiation; research in biological diversity. 72:387–395.

Wells, J.M., and F.M. Watt. 2018. Diverse mechanisms for endogenous regeneration and repair in mammalian organs. Nature. 557:322–328.

Whitbread, L.A., and B.C. Powell. 1998. Expression of the intermediate filament keratin gene, K15, in the basal cell layers of epithelia and the hair follicle. Experimental cell research. 244:448–459.

Yuan, Y., N. Salinas Parra, Q. Chen, and R. Iglesias-Bartolome. 2022. Oncogenic Hedgehog-Smoothened Signaling Depends on YAP1‒TAZ/TEAD Transcription to Restrain Differentiation in Basal Cell Carcinoma. J Invest Dermatol. 142:65–76 e67.

Zhan, Q., S. Signoretti, D. Whitaker-Menezes, T.M. Friedman, R. Korngold, and G.F. Murphy. 2007. Cytokeratin15-positive basal epithelial cells targeted in graft-versus-host disease express a constitutive antiapoptotic phenotype. J Invest Dermatol. 127:106–115.

Zhang, H., H.A. Pasolli, and E. Fuchs. 2011. Yes-associated protein (YAP) transcriptional coactivator functions in balancing growth and differentiation in skin. Proc Natl Acad Sci U S A. 108:2270–2275.

