## Supplementary material for "Keratin 15 promotes a progenitor cell state in basal keratinocytes of skin epidermis": Redmond et al. Suppl. FIGs 1-5 & legends

**List of elements:**

**Supplemental Figure 1** (complement to Figure 2).
Analysis of human trunk skin using scRNAseq.

**Supplemental Figure 2** (complement to Figure 2).
Relationships between *KRT14, KRT15 and KRT10* in human foreskin.

**Supplemental Figure 3** (complement to Figure 2).
Relationships between *Krt14, Krt15 and Krt10* in mouse back skin.

**Supplemental Figure 4** (complement to Figure 6).
Keratin and 14-3-3 interaction analyses.

**Supplemental Figure 5** (complement to Figure 7).
*Krt14*$^{C373A/null}$ protein and transcriptional analyses**.**

**Legends to supplemental figures:**

**Supplemental Figure 1** (complement to Figure 2).
**Analysis of human trunk skin using scRNAseq.**
**A.** UMAP of scRNAseq data collected from human trunk skin. Clusters consists of 17 keratinocyte clusters and 3 clusters matching non-keratinocyte signature genes (melanocytes, cl. 7 and 18; immune cells, cl. 17). **B,C.** Feature map plotting expression levels of (B) *KRT14* or (C) *KRT10* in the human trunk UMAP. **D.** Expression of *KRT14* (green) and *KRT10* (red) in the human trunk dataset, as described in Figure 2C. **E.** Feature map of *KRT15* expression levels in human trunk dataset presented as UMAP form. **F,G.** Expression of S phase composite score relative to (F) *KRT15* and (G) *KRT14* levels across keratinocytes in human trunk epidermis. **H.** Expression of G2M composite score relative to *KRT10* levels across keratinocytes in human trunk epidermis.

**Supplemental Figure 2** (complement to Figure 2).
**Relationships between *KRT14, KRT15 and KRT10* in human foreskin.**
**A,B.** UMAP of human foreskin highlighting (A) *KRT15* (green) vs. *KRT14* levels (red) and (B) *KRT15* (green) vs. *KRT10* levels (red) across Seurat clusters. **C.** Single cell expression of *KRT15* mapped against *KRT14* levels in 24,768 keratinocytes in the foreskin dataset. **D.**

Distribution of *KRT14* and *KRT15* expression across all keratinocytes in dataset. **E.** Distribution of *KRT15, KRT14* and *KRT10* expression in cells exhibiting increasing levels of 'Basal Score' signature genes *(composite scores defined in Methods).* **F.** Analysis of average *KRT14* (red, circle), *KRT15* (green, square) and *KRT10* (blue, triangle) expression level across nine Seurat clusters matching keratinocyte signatures in the foreskin dataset. Clusters are sorted on the X axis from those showing highest 'basal score' to highest 'differentiation score'. Seurat cluster matching the highest level of G2M composite score is highlighted. **G.** Expression of G2M composite score relative to *KRT14* and *KRT15* levels across keratinocytes in the foreskin. **H.** Comparison of linear pairwise expression between type I and type II keratins *KRT14/KRT5* (red, $R^2$=0.86), *KRT10/KRT1* (blue, $R^2$=0.92), and *KRT15/KRT5* (green, $R^2$=0.66) across all keratinocytes in the foreskin dataset. **H'.** Heatmap summarizing Pearson (r) correlations between type I and type II keratins across the dataset.

**Supplemental Figure 3** (completement to Figure 2).
**Relationships between *Krt14, Krt15 and Krt10* in mouse back skin.**
(A-B) UMAP of saline-treated, tape stripping sensitized mouse back skin highlighting *Krt15* levels (green) and (A) *Krt14* or (B) *Krt*10 levels (red) across Seurat clusters. (C) Single cell expression of *Krt15* mapped against *Krt14* levels in 4,806 keratinocytes in the mouse back skin dataset. (D) Distribution of *Krt14* and *Krt15* expression across all keratinocytes in the dataset. (E) Distribution of *Krt15, Krt14* and *Krt10* expression in cells exhibiting increasing levels of 'Basal Score' signature genes *(composite scores defined in Methods).* (F) Analysis of average keratin expression level across 7 Seurat clusters matching keratinocyte signatures in the back skin dataset. *Krt14* (red, circle), *Krt15* (green, square) and *Krt10* (blue, triangle) assessed per cluster. Clusters are sorted on the X axis from those showing highest 'basal score' to highest 'differentiation score'. Seurat cluster matching the highest level of G2M composite score is highlighted. (G) Expression of G2M composite score relative to *Krt14* and *Krt15* levels across keratinocytes in mouse sensitized back skin. (H) Comparison of linear pairwise expression between type I and type II keratins *Krt14/Krt5* (red, $R^2$=0.65), *Krt10/Krt1* (blue, $R^2$=0.19), and *Krt15/Krt5* (green, $R^2$=0.50) across all keratinocytes in the mouse back skin dataset. (H') Heatmap summarizing Pearson (r) correlations between type I and type II keratins across the dataset.

**Supplemental Figure 4** (complement to Figure 6).
**Analyses of keratin and 14-3-3 protein interactions**
Lollipop plots displaying the *14-3-3Pred* "Consensus Score" for all serines and threonines in **A.** human *KRT14*; **A'.** mouse *Krt14*; **B.** human *KRT5*; **B'.** mouse *Krt5*; **C.** human *KRT15*; **C'.** mouse *Krt15*. Lollipop height and color both scale to Consensus Score magnitude. The two residues with the largest positive Consensus Score are labeled. **D.** Lollipop plot illustrates identified phosphorylation sites on K14 protein exhibiting a localization score >0.6 in human NTERT keratinocytes in culture. The relative occupancy percentage for each phosphorylation site was calculated by dividing the phosphopeptide's intensity by total intensity of unique keratin 14 peptides. This calculation, shown here as relative phosphorylation site occupancy (%), was performed only for phosphopeptides that are unique to keratin 14 without miscleavages (S44, S281, S435, S437). **D'.** Spectra of the phosphopeptides corresponding to the S39 (left) and S44

(right) sites. "Phos" conveys phosphorylation, and "CAM" stands for carbamidomethyl. **E.** Representative micrographs of YAP1-14-3-3σ proximity ligation assay performed on transfected HeLa cells. Cells were transfected with untagged keratin 5 and WT and mutant K14 or K15. Merge panels show PLA punctae (red), EGFP-tagged keratin autofluorescence (green), and DAPI counterstain (blue). Scale bar = 10 µm.

**Supplemental Figure 5** (complement to Figure 7).
**Molecular analyses of *Krt14*$^{C373A/null}$ transgenic mouse skin**.
**A.** K14 and **A'.** K15 dot blots of whole tail skin lysate from 8-week old male *Krt14*$^{C373A/WT}$ and *Krt14*$^{C373A/null}$ littermates. For K14 immunoblotting, 7.5 µg, 5 µg, 2.5 µg, 1 µg, and 0.5 µg of lysate were loaded onto membrane. For K15 immunoblotting, 5 µg, 2.5 µg, 1 µg, 0.75 µg, and 0.5 µg of lysate were loaded onto membrane. Histone H3 was utilized as a loading control. Mean of K14 and K15 dot blot mean gray value normalized to Histone H3. Comparisons made using Mann-Whitney tests. *** P = 0.0006. ** P = 0.0025. **B.** Transcription of a YAP1 target gene, *Cyr61*, as measured via RT-qPCR in whole tail skin lysate from 8-week old male and female *Krt14*$^{C373A/WT}$ and *Krt14*$^{C373A/null}$ littermates. Dots represent 3 biological replicates. Comparisons made using Mann-Whitney tests. * P <0.005.

### Redmond et al. - Supplemental Figure 1

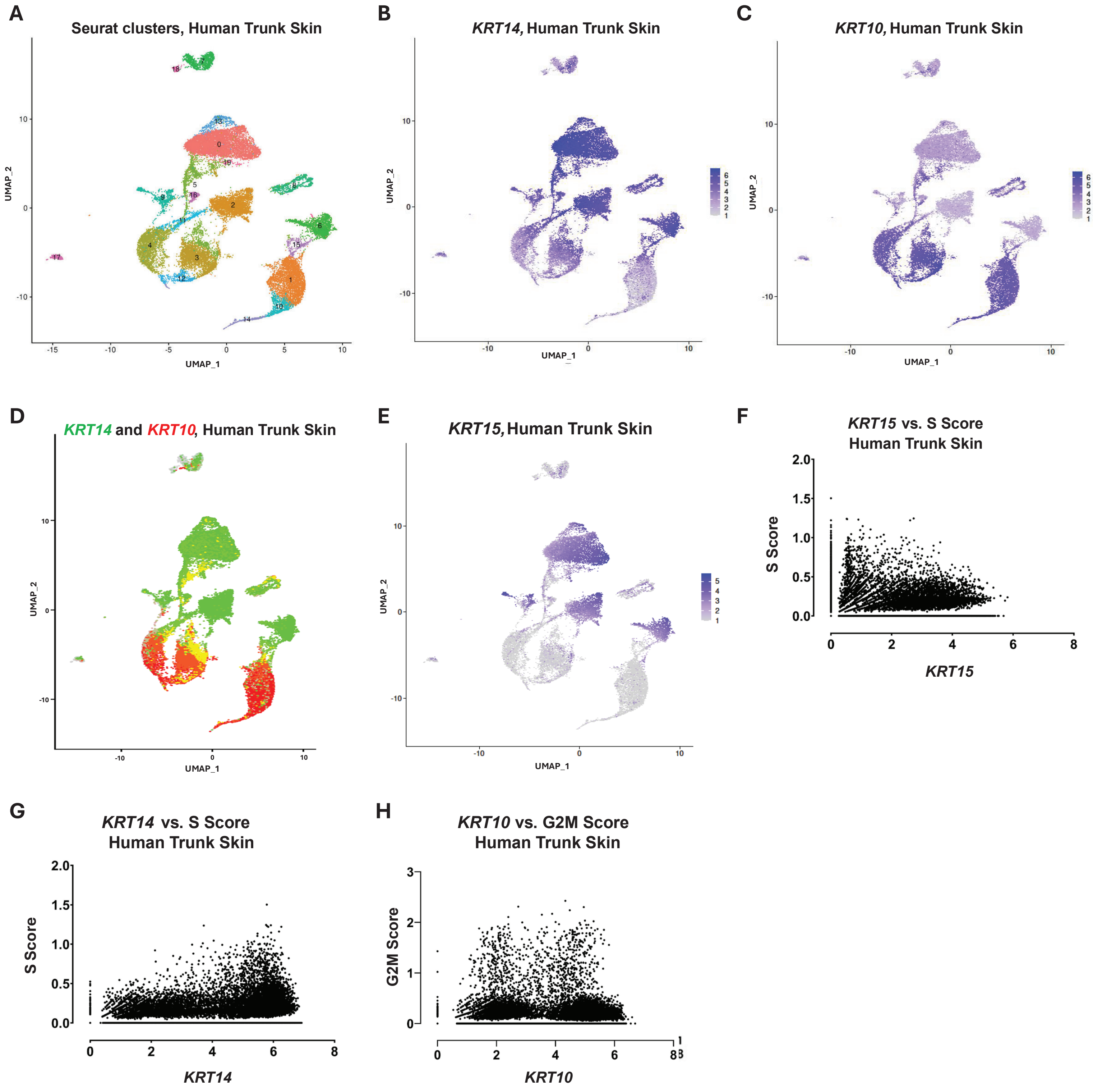

Redmond et al. - Supplemental Figure 2

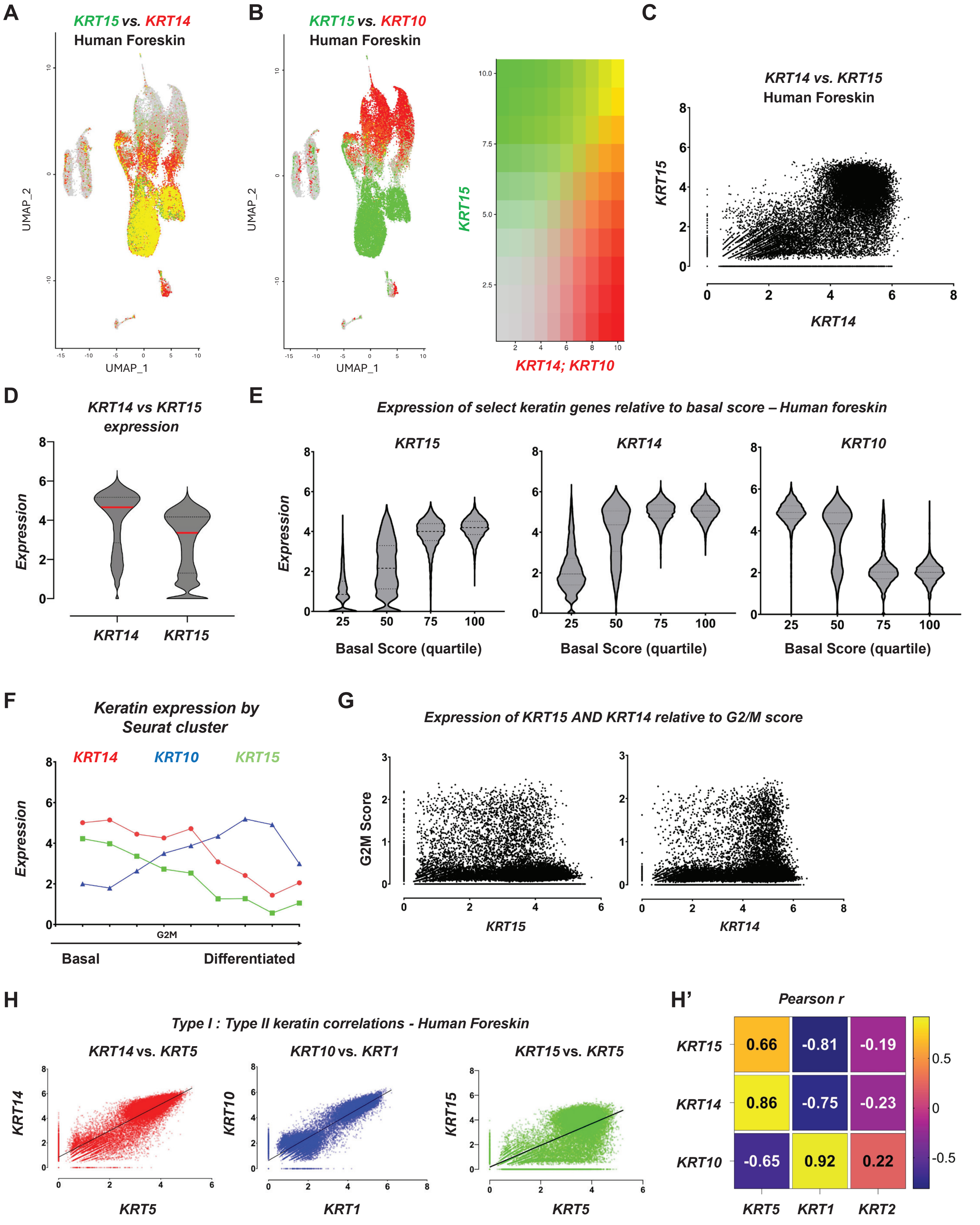

### Redmond et al. - Supplemental Figure 3

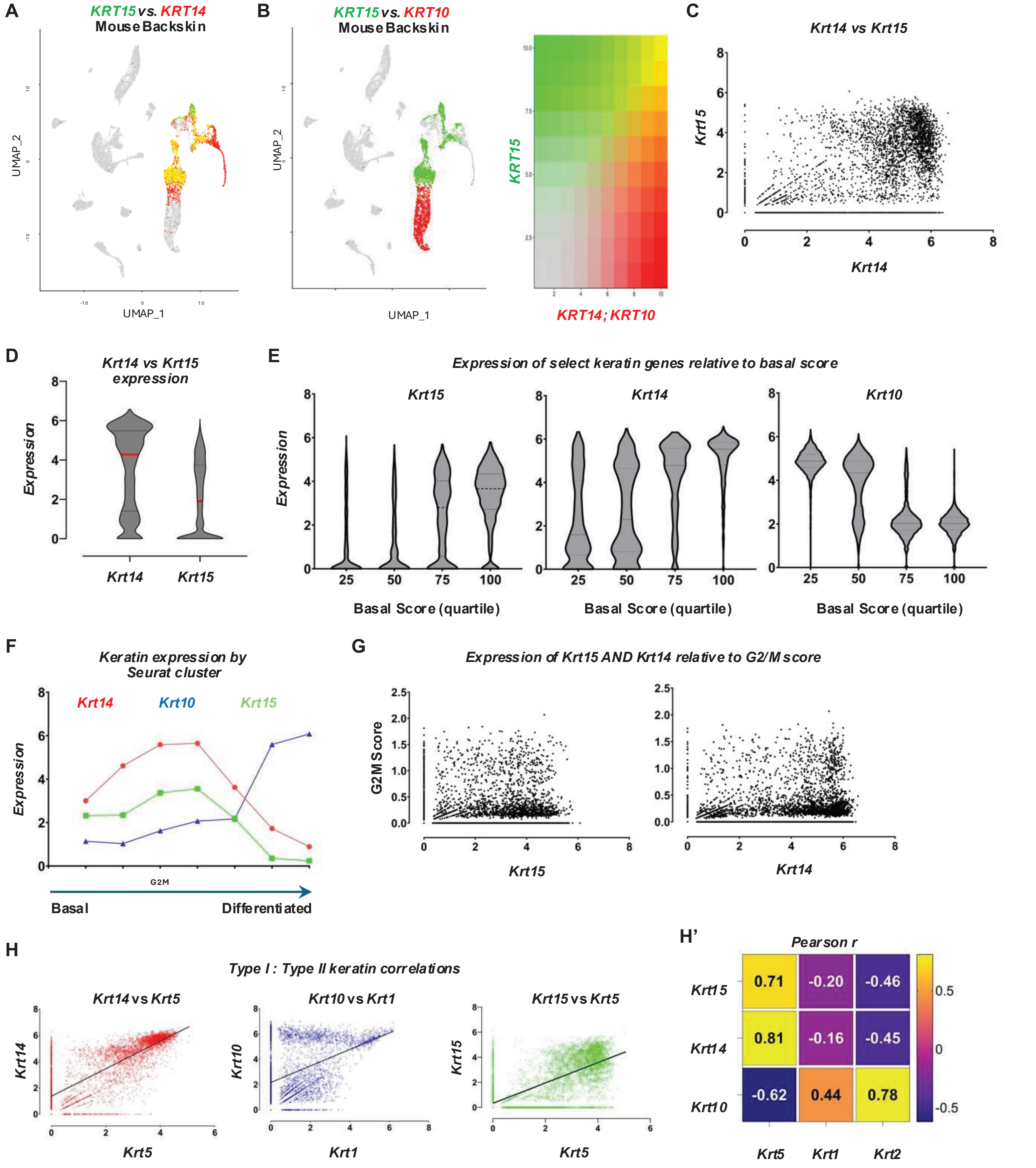

**Redmond et al. - Supplemental Figure 4**

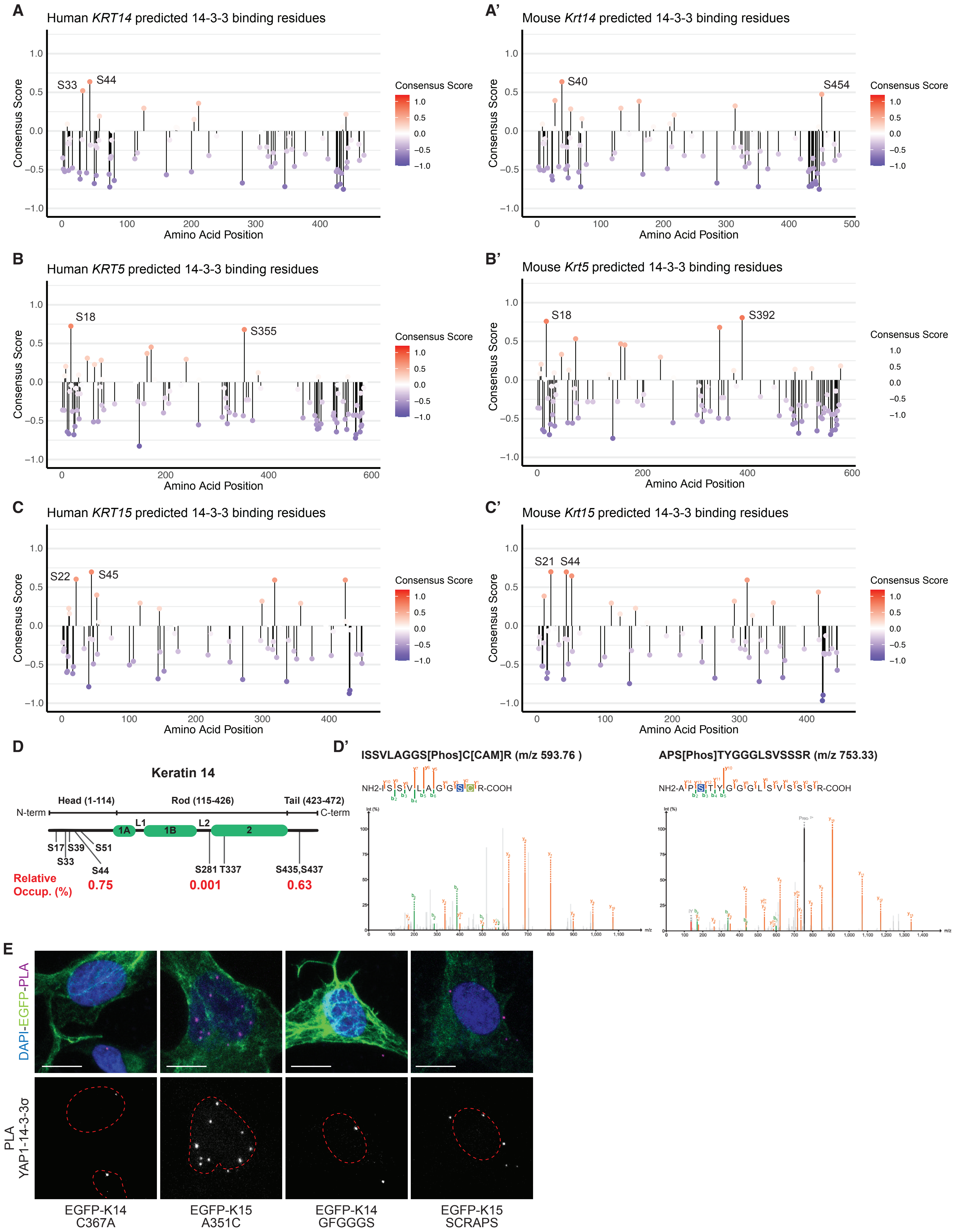

### Redmond et al. - Supplemental Figure 5

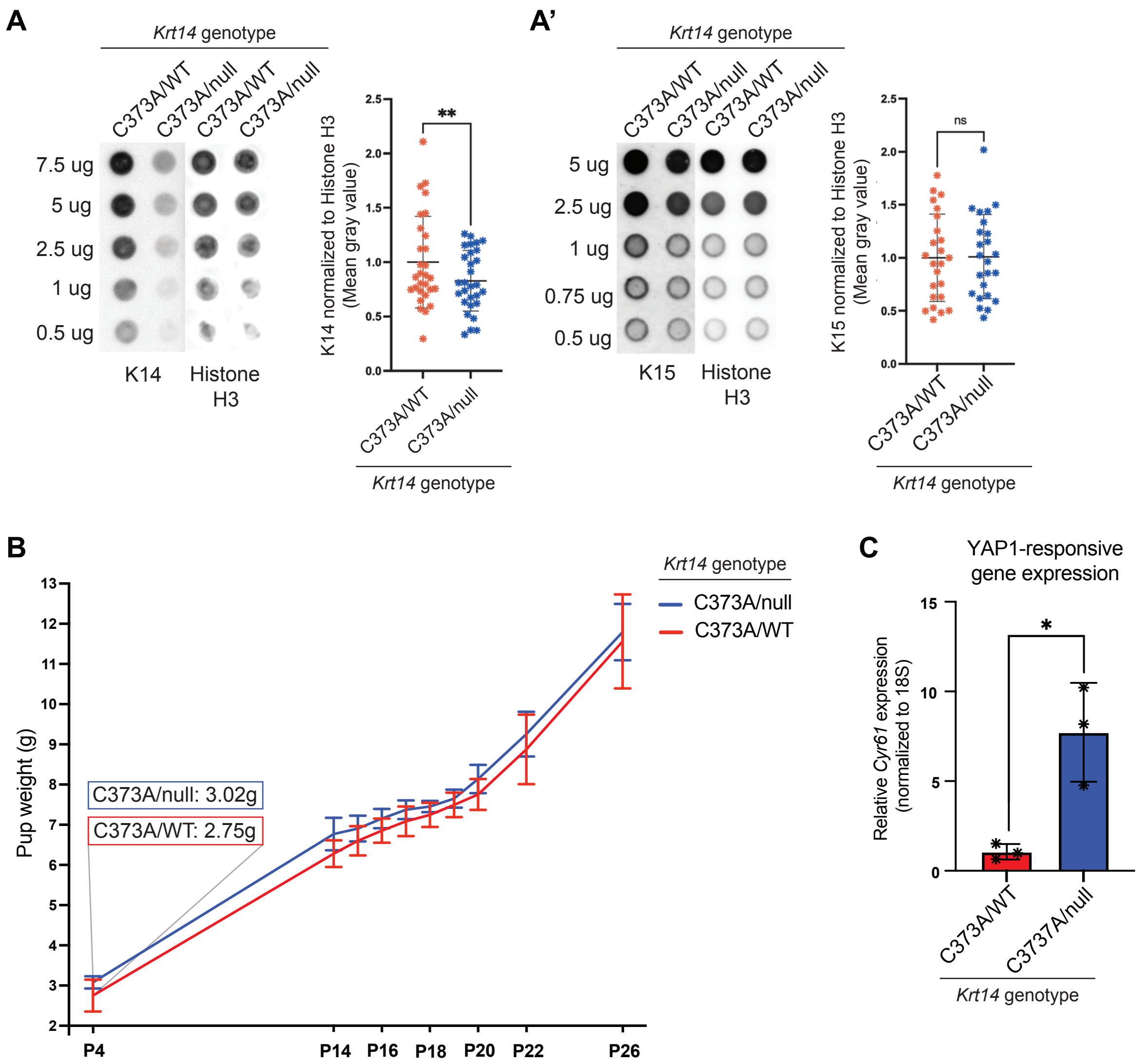
